# Single molecule tracking reveals the role of transitory dynamics of nucleoid-associated protein HU in organizing the bacterial chromosome

**DOI:** 10.1101/2019.12.31.725226

**Authors:** Kelsey Bettridge, Subhash Verma, Xiaoli Weng, Sankar Adhya, Jie Xiao

## Abstract

HU is the most conserved nucleoid-associated protein in eubacteria and has been implicated as a key player in global chromosome organization. The mechanism of HU-mediated nucleoid organization, however, remains poorly understood. Using single molecule tracking coupled with genetic manipulations, we characterized the dynamics of HU in live *Escherichia coli* cells. We found that native HU dimers bind and unbind chromosomal DNAs weakly and transitorily across the entire nucleoid volume but remain nucleoid-localized, reminiscent of random diffusion in a liquid phase-separated, membrane-less “macro-compartment” distinct from the remaining cytosol. Mutating three key surface lysine residues of HU nearly entirely abolished the weak and transitory interactions of HU with DNA and led to severe cell growth and DNA segregation defects, suggesting the importance of HU’s interactions with chromosomal DNA mediated by the positively charged surface. A conserved proline residue important for recognizing bent and cruciform DNAs such as that in recombination intermediates, similarly abolished HU’s rapid and transitory DNA interaction dynamics but had little impact on its apparent binding stability with nonspecific chromosomal DNAs. Interestingly, the proline residue appeared to be important for HUαβ dimer formation as mutating this residue makes HUαβ behave similarly to HUα_2_ dimers. Finally, we find that while prior evidence has found HU capable of depositing nucleoid-associated noncoding RNAs onto cruciform DNA structures, deletion of these specific naRNAs or inhibition of global transcription had a relatively minor effect on HU dynamics irrespective altered nucleoid compaction. Our results suggest a model of chromosome organization mediated by weak, transient interactions of HU, a substantial deviation from nucleoid-like proteins such as histones. Such collective sum of the numerous weak, transitory binding events of HU with nonspecific chromosome DNAs could generates a “force” to maintain a dynamic, fluid nucleoid with enough flexibility to rapidly facilitate global topological processes such as replication or nucleoid segregation.

## Introduction

The *E. coli* genome must condense over a thousand-fold to be packed into a cell of a few micrometers in size. This condensed genome, together with its associated proteins and RNAs, is termed the nucleoid and organized into six spatially isolated macrodomains (MD)^1–5^. The precise molecular mechanism of how this nucleoid organization is achieved is not well understood, but many key players have been identified. A group of proteins known as nucleoid-associated proteins (NAPs), such as H-NS, IHF, Fis, and HU have been shown to play important roles (reviewed in ^3^). Among these NAPS, HU is the most conserved across eubacteria and the one of the most abundant in *E. coli* (∼30,000 copies per cell during middle exponential phase growth^6, 7^). *E. coli* HU has two subunits of ∼9 kDa each, α and β, which form either HUα_2_ homodimers or HUαβ heterodimers depending on the growth phase (HUβ_2_ homodimers were negligible under log-phase growth)^7^. Monomers of HU are not detectable^6, 7^. HU was originally proposed to be the bacterial equivalent to eukaryotic histones due to its small size, lack of sequence specificity, and its *in vitro* ability to generate condensed plasmid DNA^8^ and to induce negative supercoils in relaxed plasmid DNA in the presence of topoisomerase I^9^. Additionally, HU impacts numerous cellular processes including replication^10, 11^, transcription^12^ and translation^13^, and has been implicated in pathogenesis^14^, likely through its role in organizing the chromosomal structure.

Despite its importance, how HU mediates nucleoid organization has remained elusive. HU binds specifically with high affinity to distorted DNAs such as kinked or cruciform DNAs that are recombination intermediates^15^, but for bulk chromosomal double-stranded (ds) DNAs the binding is weak and nonspecific^16^. How such weak and nonspecific binding impacts nucleoid organization is unclear, but many models have been proposed. One early model implicated HU’s role in the maintenance of the supercoiling state of the nucleoid, as deletion of HU resulted in sensitivity to UV radiation^17^, antibiotic novobiocin (inhibiting gyrase)^18^, and cold shock^19, 20^. Suppressor mutations in gyrase^18^ and increased expression of topoisomerase I^21^ could compensate the loss of HU. Other effects include altered global transcription^12^, defects in replication, and chromosome mis-segregation^20^. However, the supercoiling maintenance model was challenged by a more recent study probing the global supercoiling state using psoralen crosslinking, and demonstrated HU maintains the supercoiling state only during the stationary phase^22^. An *in vitro* study using single molecule optical tweezers found that at higher concentrations, HU could change the persistence length of DNA^23^, suggesting HU could alter the physical properties of the DNA. Consistent with this suggestion, loss of HU was shown to affect small loop formation *in vivo*^24^ and HU is required for DNA loop-mediated repression of the *gal* operon^25^. Furthermore, HMGB1, the eukaryotic analog of HU, significantly increases circularization of small (<200 bp) DNA fragments^26^ and could partially replace HU in a looping assay in an HU deletion background^27^.

More recently, a study demonstrated that HU binds to RNAs^13, 28, 29^, and that in the presence of HU, post-processed small non-coding RNAs (renamed as nucleoid-associated RNAs, naRNAs) from repetitive extragenic palindromic (*REP*) elements^29–31^ condensed plasmids *in vitro* and promoted chromosomal contacts between various *REP* elements *in vivo*^30^. These observations lead to a catalytic chaperone model, in which HU binds both cruciform DNA and naRNAs to bridge DNA-DNA contacts and mediate chromosomal organization^31^. Another model, based on the compatible interfaces of HUα_2_ and HUαβ dimers found in the crystal structures of HU bound to non-specific dsDNA, suggests that HU compacts the nucleoid through enhanced DNA bundling by oligomerizing on linear dsDNAs^32^. However, HU oligomerization has not been directly observed *in vivo*. Each of these models involves a distinct physical mechanism (maintenance of supercoiling, RNA chaperon, or oligomerization of HU) that was deduced from genetic and *in vitro* biochemical studies. How HU interacts with chromosomal DNA *in vivo* and how such interactions compact the nucleoid are unknown.

In this work, we examined the *in vivo* interaction dynamics of HU with chromosomal DNAs using single-molecule tracking (SMT) under conditions of wild-type (WT) and perturbed DNA binding, hetero-dimerization or RNA binding of HU. Our results directly demonstrate that HU interacts with chromosomal DNAs weakly and transitorily but remains highly nucleoid-localized. These interactions are likely mediated through the positive surface charge of the HU dimer, as mutations of three lysine residues on the surface lead to a loss of HU nonspecific DNA interaction dynamics. Mutation of a conserved proline residue thought to be important for DNA bending showed a less severe change in HU dynamics and cell phenotype and was remarkably similar to HUα2 dimer behavior, suggesting the proline residue is important for HUαβ heterodimer function or formation. Loss of specific naRNAs or global downregulation of transcription altered the nucleoid morphology, but did not significantly impact HU dynamics or localization, suggesting HU does not play a stable architectural role in chromosome organization. Ultimately, our results implicate a model where the major function of HU in chromosome organization is through maintaining the viscoelastic properties of the nucleoid through numerous weak, nonspecific interactions with chromosomal DNAs to enable facilitation of global topological processes such as replication, gene expression, and nucleoid segregation.

## Results

### HU-PAmCherry exhibited two rapidly switching diffusive states

To probe the dynamics of HU in the nucleoid, we used an *E. coli* strain in which the chromosomal copy of *hupA* gene encoding for the α subunit of HU was replaced by a 3’ tagged photoactivatible fluorescent protein fusion gene *hupA-PAmCherry*^33^. We verified that the fusion protein HUα-PAmCherry was expressed at the expected full length as the sole cellular source of HUα (Figure S1A) and supported wild-type (WT)-like growth (Figures S1B and S1C). Furthermore, HUα-PAmCherry supported HU-dependent mini-P1 plasmid replication and Mu phage growth in the Δ*hupB* background at levels indistinguishable from those of the WT parental strain MG1655^29, 30^ (Figures S1D and S1E). These results suggested HUα-PAmCherry could replace the endogenous HUα for its function.

To perform single molecule tracking on HUα-PAmCherry, we imaged mid log-phase live *E. coli* cells grown in EZ rich defined media (EZRDM) at room temperature (24 °C) on agarose gel pads. We activated individual HUα-PAmCherry molecules using a low level of 405 nm UV light and tracked their cellular locations with a frame rate of ∼150 Hz (Δ*t* = 6.74ms) with a laser excitation at 568 nm. This imaging rate allowed us to determine the diffusion coefficients of individual molecules up to 3 μm^2^/s with high confidence. In Figure 1A, we showed a few representative trajectories superimposed with the corresponding bright-field image of the cell. Individual trajectories displayed different diffusive properties: some were confined within small regions (blue trajectories), some traversed larger cell areas (red trajectories), and some switched in between (rainbow trajectories). Plotting the apparent diffusion coefficients calculated from single-step displacements of all molecules indeed suggested that there existed a heterogeneous distribution of HUα-PAmCherry molecules with a wide range of diffusion coefficients (Figure S2A).

**Figure 1:**
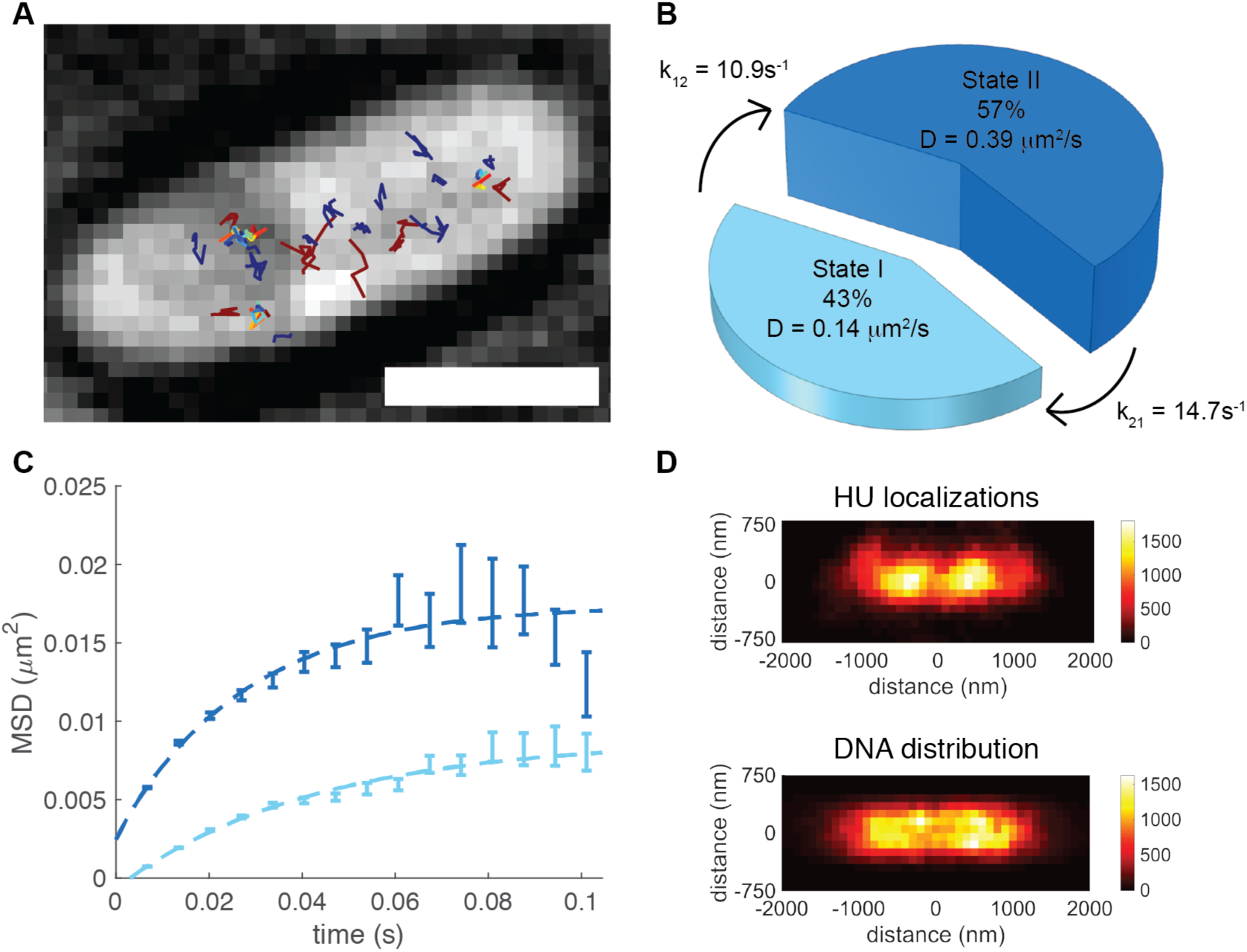
Single molecule tracking (SMT) of HUα-PAmCherry in live *E. coli* cells grown in rich media showed two diffusive states, rapid transition kinetics, and nucleoid-like localizations. (A) Representative SMT trajectories superimposed on top of the corresponding brightfield image of the cell (gray). Blue trajectories represented State I and red State II. Molecules transitioned between the two states were colored in rainbow. Scale bar = 1 μm. (B) Two diffusive states of HUα-PAmCherry molecules with respective population percentages (size of the pie piece), transition rates, and diffusion coefficients (height of pie piece) as identified by the HMM. (C) Mean squared displacement (MSD) plots of State I (light blue) and State II (dark blue) HUα-PAmCherry trajectories as a function of time; the plateau signified confined diffusion of molecules in both States. (D) Two-dimensional (2D) histogram of all cellular HUα-PAmCherry localizations from SMT (top) and aggregated nucleoid morphology from SIM (structured illumination microscopy) imaging (bottom). The pixel size of both was 100 × 100 nm. The top color bar indicated the number of localizations used in each bin for HUα-PAmCherry (total 59,432 localizations). The bottom color bar indicated the normalized fluorescence level (in arbitrary unit) of the nucleoid-intercalating dye Hoechst 33342 (total 20 fluorescence images).

To analyze different populations quantitatively, we used a Bayesian-based hidden Markov Model (HMM)^34^, which robustly determines the optimal number of diffusive states represented in the data, the diffusion coefficients of those states, the percentage of molecules in each state, and the apparent transition rates between states. We found that the data fit best to a two-state model (Figure 1B, Table 1), with diffusion coefficients of State I *D*_1_ = 0.14 ± 0.004 μm^2^/s, and State II *D*_2_ = 0.39 ± 0.006 μm^2^/s, (μ ± s.e.m., n = 60,432 displacements). The percentages of the HU molecules inState I and II were ∼45% and ∼55% respectively. Using several other classic methods^35^, we obtained the same two states with essentially identical diffusion coefficients and population percentages (Figures S2A, S3A and S4A; Table S3; Supplemental Note 1). The presence of two apparent diffusive states likely reflects the binding and unbinding interactions of HUα-PAmCherry to chromosomal DNAs in the nucleoid^13, 15, 22^. Furthermore, using HMM, we found that HUα-PAmCherry molecules switched between the slow diffusing State I and the fast diffusing State II with apparent rates of *k_12_* = 10.9 s^-1^ and *k_21_* = 14.9 s^-1^ respectively (Figure 1B). The dwell time of States I and II were at 100 ± 15 ms and 74 ± 7 ms, respectively (Table 1). The fast switching kinetics of HU-PAmCherry between the two states indicate that these interactions were rapid and transitory.

**Table 1:** Diffusion parameters extracted from single-moleucle tracking trajectorioes of Huα-PAmCherry under different experimental conditions using the Hidden Markov Model (HMM). All values expressed as µ ± s.e.m.

|  | State I |  |  | State II |  |  | n<br>(displacements) |
| --- | --- | --- | --- | --- | --- | --- | --- |
| | $D_1$ ($\mu\text{m}^2/\text{s}$) | Occupation (%) | Dwell time (ms) | $D_2$ ($\mu\text{m}^2/\text{s}$) | Occupation (%) | Dwell time (ms) | |
| Huα-PAmCherry | $0.14 \pm 0.004$ | $45 \pm 2$ | $100 \pm 15$ | $0.38 \pm 0.006$ | $55 \pm 2$ | $74 \pm 7$ | 60,432 |
| Huα(*triKA*)-PAmCherry | $0.18 \pm 0.01$ | $5 \pm 0.2$ | $107 \pm 23$ | $1.08 \pm 0.04$ | $95 \pm 0.2$ | $852 \pm 142$ | 193,678 |
| Huα(*P63A*)-PAmCherry | $0.25 \pm 0.01$ | $32 \pm 1$ | $129 \pm 15$ | $0.84 \pm 0.01$ | $68 \pm 1$ | $114 \pm 8$ | 90,406 |
| Huα-PAmCherry + chlor | $0.13 \pm 0.003$ | $40 \pm 1$ | $80 \pm 7$ | $0.35 \pm 0.003$ | $60 \pm 1$ | $70 \pm 4$ | 108,846 |
| Huα-PAmCherry + rif | $0.14 \pm 0.003$ | $22 \pm 2$ | $80 \pm 8$ | $0.39 \pm 0.001$ | $78 \pm 2$ | $100 \pm 7$ | 137,019 |
| Huα-PAmCherry + ΔRep325 | $0.13 \pm 0.004$ | $30 \pm 1$ | $72 \pm 9$ | $0.36 \pm 0.003$ | $70 \pm 1$ | $96 \pm 9$ | 43,413 |
| Huα-PAmCherry + Δ*hupB* | $0.24 \pm 0.01$ | $38 \pm 2$ | $168 \pm 43$ | $0.88 \pm 0.02$ | $62 \pm 2$ | $155 \pm 29$ | 22,449 |
| Huα(*triKA*)-PAmCherry + Δ*hupB* | $0.41 \pm 0.01$ | $42 \pm 0.8$ | $196 \pm 23$ | $2.42 \pm 0.03$ | $58 \pm 0.8$ | $84 \pm 6$ | 58,867 |
| Huα(*P63A*)-PAmCherry + Δ*hupB* | $0.29 \pm 0.02$ | $30 \pm 1$ | $100 \pm 8$ | $1.02 \pm 0.05$ | $70 \pm 1$ | $110 \pm 6$ | 132,631 |

### The two diffusive states reflected HU-PAmCherry’s transient and nonspecific binding with chromosomal DNA

Based on the values of the apparent diffusion coefficients, we reasoned that the slow diffusive State I (*D*_1_ = 0.14 ± 0.004 μm^2^/s) likely represented HUα-PAmCherry molecules bound to chromosomal DNA, whereas the fast diffusive State II molecules (*D*_2_ = 0.39 ± 0.006 μm^2^/s) the unbound population diffusing in the nucleoid. Note that the dwell times of both states were significantly shorter than what would be expected for the specific binding of most DNA binding proteins^36^ (Table 1), or the known nanomolar affinity of HU to nicked, cruciform, and kinked DNA structures^15, 28^, but were consistent with the weak binding nature of HU to the bulk chromosomal dsDNA^16^. Supporting this possibility, we found that the mean-squared displacement (MSD) curve of State I molecules plateaued at a level significantly lower than to that of State II molecules at relatively long time scales (> 0.1 s, Figure 1C), suggesting that State I molecules experienced more restricted diffusion than State II molecules (Figure 1C). Using a confined diffusion model assuming a finite circular boundary (Supplemental Note 2), we estimated the diameter of the confinement zone of single State I molecules at ∼230 nm, significantly larger than that determined from stationary HUα-PAmCherry molecules bound to DNA in chemically fixed cells (Figure S5, ∼110 nm). It is possible that State I HUα-PAmCherry molecules did not remain bound to DNA during the ∼100 ms dwell time but “hopped” on adjacent sequences transiently, or that they remained bound, but the bound chromosomal DNA had intrinsic motions on the same time scale in live cells.

To differentiate these two possibilities, we tracked the diffusion of a chromosomal DNA segment labeled with six *tetO* operator sites (*tetO*^6^) tightly bound by TetR-mCherry fusion protein molecules under the same cell growth and imaging conditions as HUα-PAmCherry (Figure S6A). Note that compared to other fluorescent repressor-operator systems (FROS) using tandem arrays of hundreds of DNA binding sites^37^, the small footprint of the *tetO*^6^ site (∼200 bp) enabled us to pinpoint the position of the labeled chromosomal DNA with high accuracy and negligible perturbations^38^. We observed that TetR-PAmCherry formed distinct fluorescent spots in cells (Figure S6A), consistent with its specific DNA binding property. In contrast, the cellular distribution of HU-PAmCherry mimicked that of the nucleoid (Figure 1D), indicating that the binding sites of HU are distributed throughout the chromosome. The average apparent diffusion coefficient *D*_app_ of TetR-PAmCherry molecules (0.09 ± 0.04 μm^2^/s, μ ± s.e., n = 1808 displacements, Figure S6B) and the corresponding confinement zone (∼200 nm, Figure S6C) were nearly identical to those of State I HUα-PAmCherry molecules. Note that the *D*_app_ of ∼ 0.09 was also comparable to what was previously reported on labeled *E. coli* chromosomal foci (∼ 0.1 μm^2^/s)^39^. These results suggest that State I HUα-PAmCherry molecules most likely represented the DNA-bound population even though the binding was short-lived, and that the chromosomal DNA segments were intrinsically dynamic.

For State II HUα-PAmCherry molecules, the *D*_app_ at ∼0.4 μm^2^/s was one order of magnitude smaller than what would be expected from a freely diffusing, non-DNA-interacting molecule of similar size in the *E. coli* cytoplasm (1 to 10 μm^2^/s)^40^, but comparable to those of the unbound population of many other DNA-binding proteins in *E. coli*, such as LacI^41^, RNA polymerase^42^, and UvrA^43^. Furthermore, the apparent diffusion domain diameter of State II HUα-PAmCherry molecules was ∼ 300 nm, smaller than the size of nucleoid, which we estimated at ∼550 nm (FWHM) along the short axis of cell based on structured illumination imaging under the same growth condition (Figure 1D and S7). These results suggest that when dissociated from DNA, a HUα-PAmCherry molecule likely did not diffuse far before it encountered another nonspecific binding site, which was effectively any other chromosomal DNA sequences nearby. These frequent interactions are not resolvable under our imaging time resolution (frame rate of 150 Hz, Δ*t* = 6.74ms) and hence the observed apparent diffusion coefficient reflects the overall, averaged nonspecific interactions of HU with chromosomal DNA. Less frequent interactions would speed up the diffusion and lead to larger apparent diffusion coefficient and *vice versa*. The slowed diffusion of HU and other DNA binding proteins in nucleoid due to frequent, transitory DNA interactions is reminiscent of random diffusion of molecules in liquid condensates, suggesting that the nucleoid is likely organized into a membrane-less macro-compartment with distinct viscosity properties from the cytosol{refs}.

Taken together, SMT of HU-PAmCherry revealed that HU binds to chromosomal DNAs across the nucleoid transitorily and nonspecifically. Using the apparent on-rate (*k*_21_ from fast state II to slow state I) and off-rate (*k*_12_ from slow state I to fast state II) measured from SMT and an average concentration of nonspecific chromosomal DNA binding sites at ∼ 5 mM under our growth condition (i.e. average 2 chromosomes per cell, with each chromosome having ∼4 × 10^6^ nucleotides, representing a total number of ∼ 8 × 10^6^ nonspecific binding sites in a total volume of ∼ 2 fL), we estimated that *in vivo* the nonspecific binding affinity *K*_d_ of HUα-PAmCherry to chromosomal DNA was at ∼ 4 mM with the association rate constant at ∼2 × 10^3^ M^-1^s^-1^. Both constants were two to three orders of magnitude lower than other specific DNA or RNA binding proteins^44, 45^. These results suggested that HU primarily interacts with chromosomal DNA in a weak and transitory manner^46, 47^.

### Surface lysine and proline residues have differential roles in HU’s chromosomal DNA interaction dynamics and binding affinities

Previous crystallographic studies have identified a set of residues on HU that are important to its DNA interactions (Figure S8). Three positively-charged lysine residues on the side surface of HU were shown to form electrostatic interactions with the negatively-charged backbone of linear dsDNA to stabilize its nonspecific binding^32^. A nearly universally conserved proline residue (P63) located in the β-branch arm of each HU subunit may intercalate into the minor groove of DNA to stabilize specific DNA conformations such as loops, bents and supercoils^24, 32, 48–50^ (Figure S8). Therefore, to further examine whether these residues contributed differentially to the transitory binding dynamics of HUα-PAmCherry, we constructed two mutant fusions, one with the three lysines on the surface of HUα mutated to alanines (K3A K18A K83A, referred to as triKA) and one with the conserved proline residue mutated to alanie (P63A). We then replaced the chromosomal *hupA* gene with the *hupA(triKA)-PAmCherry* or *hupA(P63A)-PAmcherry* fusion gene and investigated the corresponding binding dynamics of the mutant fusions.

We first performed SMT on HUα(triKA)-PAmCherry using the same growth and imaging conditions as WT HUα-PAmCherry cells. We found that HUα(triKA)-PAmCherry molecules displayed dramatically altered dynamics as compared to WT HU-PAmCherry (Figure 2A and B). Nearly all HUα(triKA)-PAmCherry molecules (∼ 95%) exhibited fast, random Brownian-like diffusion (*D*_app_ = 1.08 μm^2^/s, n = 193,678 displacements from 61 cells, Figure 2A and B, dark gray), with a very minor population (∼ 5%) diffusing at an intermediate *D*_app_ = 0.24 μm^2^/s (Figure 2A and B, light gray). The significantly increased diffusion coefficient of the fast diffusion state of HUα(triKA)-PAmCherry molecules suggests that a HUα(triKA)-PAmCherry molecule would diffuse a much longer distance before it can interact with a nonspecific chromosomal DNA sequence. Furthermore, while the apparent off-rate (*k*_12_) was negligibly affected, the apparent on-rate (*k*_21_) was reduced ∼ 10-fold compared to that of WT HUα-PAmCherry, resulting in a ∼ 10-fold higher apparent *K_d_* (Figure 2A, Table 1). These observations demonstrate a near complete loss in the ability of HUα(triKA) to bind to chromosomal DNA ↓i.e. HUα(triKA) diffuses freely throughout the nucleoid with minimal interactions with chromosomal DNA. Indeed, while the cellular distribution of the fast diffusion population of HUα(triKA)-PAmCherry still maintained the shape of the nucleoid, it expanded significantly toward the cytoplasm compared to that of WT HUα-PAmCherry (Figure 2C). Thus, the electrostatic interactions between the triple lysine residues on the HUα surface and the negatively charged backbone of chromosomal DNA may be mainly responsible for HU’s weak interactions with nonspecific DNA interactions and subsequent binding.

**Figure 2:**
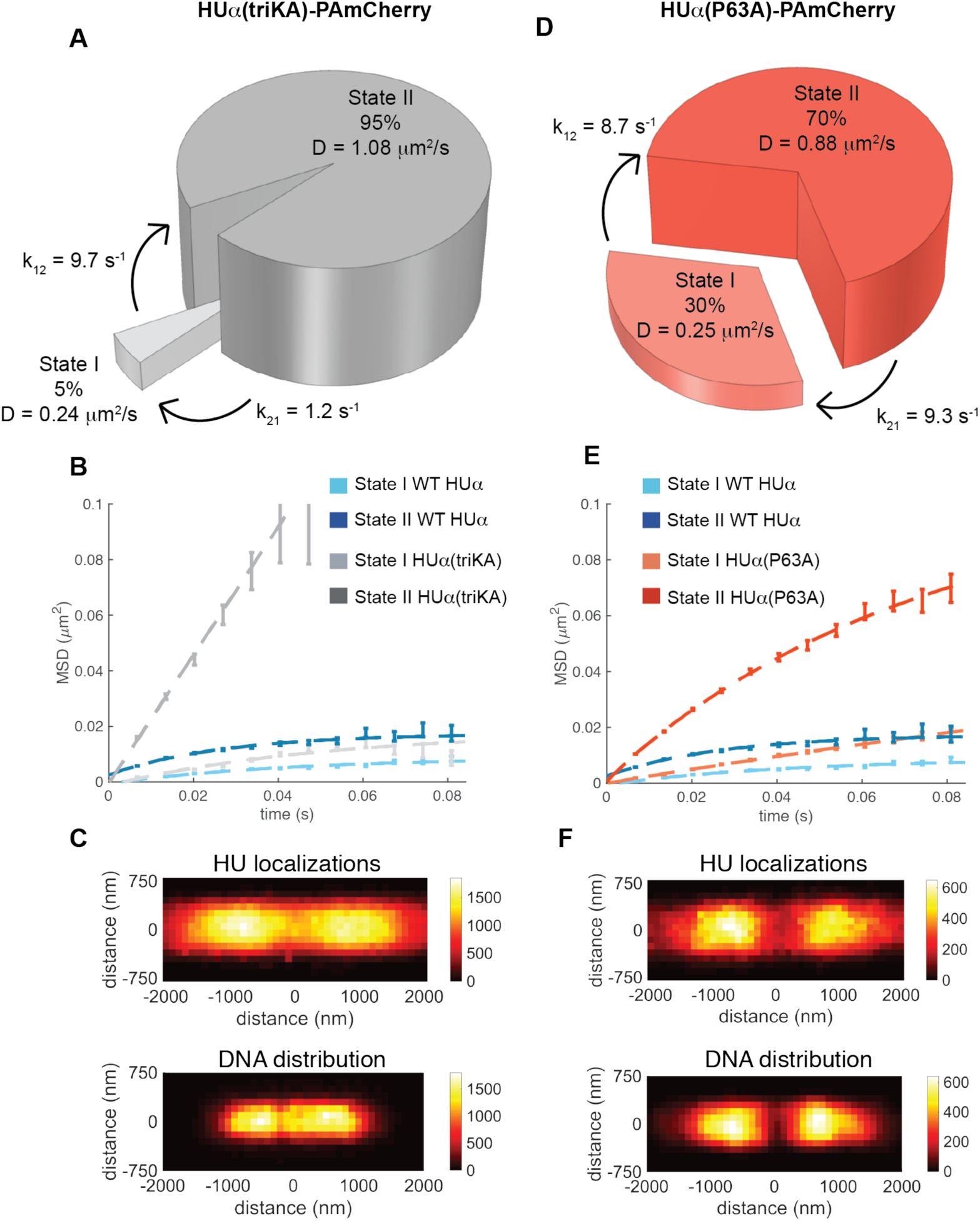
The triKA and P63A mutations affected HU dynamics differentially. (A) Two diffusive states of HUα(triKA)-PAmCherry with respective population percentages (size of pie piece), transition rates, and diffusion coefficients (height of pie piece) as identified by the HMM. (B) Mean squared displacement (MSD) plots of State I (light gray) and State II (dark gray) HUα(triKA)-PAmCherry trajectories as a function of time compared to State I and II molecules in the WT condition (light and dark blue). (C) Two-dimensional (2D) histogram of all cellular HUα(triKA)-PAmCherry localizations from SMT (top) and aggregated nucleoid morphology from SIM (structured illumination microscopy) imaging (bottom). The pixel size of both was 100 × 100 nm. The top color bar indicated the number of localizations used in each bin for HUα(triKA)-PAmCherry (total 193,678 localizations). The bottom color bar indicated the normalized fluorescence level (in arbitrary units) of the nucleoid-intercalating dye Hoechst 33342 (total 20 fluorescence images). (D) Two diffusive states of HUα(P63A)-PAmCherry with respective population percentages (size of pie piece), transition rates, and diffusion coefficients (height of pie piece) as identified by the HMM. (E) Mean squared displacement (MSD) plots of State I (light orange) and State II (dark orange) HUα(triKA)-PAmCherry trajectories as a function of time compared to State I and II molecules in the WT condition (light and dark blue). (F) Two-dimensional (2D) histogram of all cellular HUα(P63A)-PAmCherry localizations from SMT (top) and aggregated nucleoid morphology from SIM (structured illumination microscopy) imaging (bottom). The pixel size of both was 100 × 100 nm. The top color bar indicated the number of localizations used in each bin for HUα(triKA)-PAmCherry (total 90,406 localizations). The bottom color bar indicated the normalized fluorescence level (in arbitrary unit) of the nucleoid-intercalating dye Hoechst 33342 (total 20 fluorescence images).

Consistent with above observations, we observed detrimental defects of HUα(triKA) on cell growth and nucleoid morphology. Cells expressing HUα(triKA)-PAmCherry had significant wider distributions in length and width and were on average longer and skinnier than WT cells (Figure S9A, left). Most interestingly, the nucleoid of HUα(triKA) mutant cells appeared to be significantly more compact compared to that of WT HU cells (Table S7), and that the chromosomal DNA distribution in the mutant strain was no longer symmetric (Figure S7). These results suggest that the dynamic interactions of HU with chromosomal DNA and its weak binding, here likely mediated by the triple lysine residues on HUα’s surface, may play significant roles in maintaining a normal nucleoid volume and chromosome segregation.

Next, we performed SMT on HUα(P63A)-PAmCherry under the same cell growth and imaging conditions. HUα(P63A)-PAmCherry also exhibited two populations with apparent diffusion coefficients similar to HUα(triKA)-PAmCherry (*D*_app_^1^ = 0.25 ± 0.007 µm^2^/s and *D*_app_^2^ = 0.88 ± 0.008 µm^2^/s respectively, Figure 2D, Table 1, n = 90,406 displacements from 60 cells), suggesting that the P63 residue contributes significantly to the frequent interactions of HU with chromosomal DNA. However, HUα(P63A)-PAmCherry had a larger slow diffusion population (∼ 30%), and the apparent on-rate (*k_21_* = 9.3 s^-1^) was closer to that of WT cells compared to HUα(triKA)-PAmCherry. These results indicate that the P63 residue may not be involved in the binding of HU with chromosomal DNA, and hence does not impact HU’s apparent binding affinity dramatically (*K_d_*, Supplementary Table S6). Consistent with this observation, the cellular distribution of HUα(P63A)-PAmCherry remained nucleoid-centered, unlike the much more expanded distribution of HUα(triKA)-PAmCherry (Fig. 2F). Interestingly, while HUα(P63A)-PAmCherry expressing cells exhibited relatively normal growth rates and cell length distributions (Figures S1B and S9), their nucleoids expanded significantly compared to that of WT cells, opposite of what was observed in HUα(triKA)-PAmCherry expressing cells (Figure S7, Table S7). The similar effect of triKA and P63A on the diffusive property of HU indicates that both P63-mediated DNA interactions (likely on altered DNA structures such as that produced during replication, transcription and recombination^10, 11, 15, 25^) and triK-mediated DNA interactions (likely nonspecific linear dsDNA binding) contribute to the transitory interactions of HU with chromosome DNA. Their differential effects on nucleoid compaction and the apparent DNA binding affinity, however, suggest that specific-DNA structure binding of HU may be important for nucleoid compaction while linear dsDNA binding may be important for avoiding overly compacted nucleoid, which could impede DNA segregation.

### Dynamic chromosomal DNA interactions mediated by HUαβ heterodimers are important for maintaining proper nucleoid morphology and segregation

In exponentially growing WT cells, HU exists as a mixture of both HUα2 homodimers and HUαβ heterodimers, with HUβ2 dimers nearly undetectable^7^. As cells enter stationary phase, HUα2 recedes and HUαβ becomes the predominant form. It was suggested that although both HUα2 and HUαβ may function equally to organize the nucleoid by maintaining the superhelical density of the chromosomal DNA, the heterodimer may play an essential role in protecting DNA from radiation or starvation-mediated DNA damages, possibly through its differential affinity to specialized DNA structures^15^. To better understand the differential roles of the two HU dimers, we performed SMT on HUα-PAmCherry molecules in a *ΔhupB* background, in which Huα could only form homodimers of HUα2. Like WT HUα-PAmCherry, Δ*hupB* HUα-PAmCherry cells grew similarly compared to a WT parental background strain and supported HU-dependent mini-P1 and Mu phage replication (Figure S1). Despite relatively minor perturbations on cell physiology, we found that HUα_2_-PAmCherry molecules in the *ΔhupB* background diffused much faster compared to that in the WT condition (Figure 3A, B), while its apparent binding affinity to DNA (determined by the transition kinetics) was not significantly reduced (Supplementary Table S6). These observations suggest that under our growth conditions, HUαβ heterodimer is likely the major species interacting with chromosomal dsDNA transitorily, and that β subunit is minimally involved in chromosomal dsDNA binding. Importantly, the nucleoid volume of Δ*hupB* HUα-PAmCherry cells expanded substantially compared to that of WT cells (Figure S7B, Table S7), indicating that the transitory chromosomal dsDNA interaction of mediated by the HUαβ heterodimer, but not its dsDNA binding stability, is important for maintaining a compact nucleoid.

**Figure 3:**
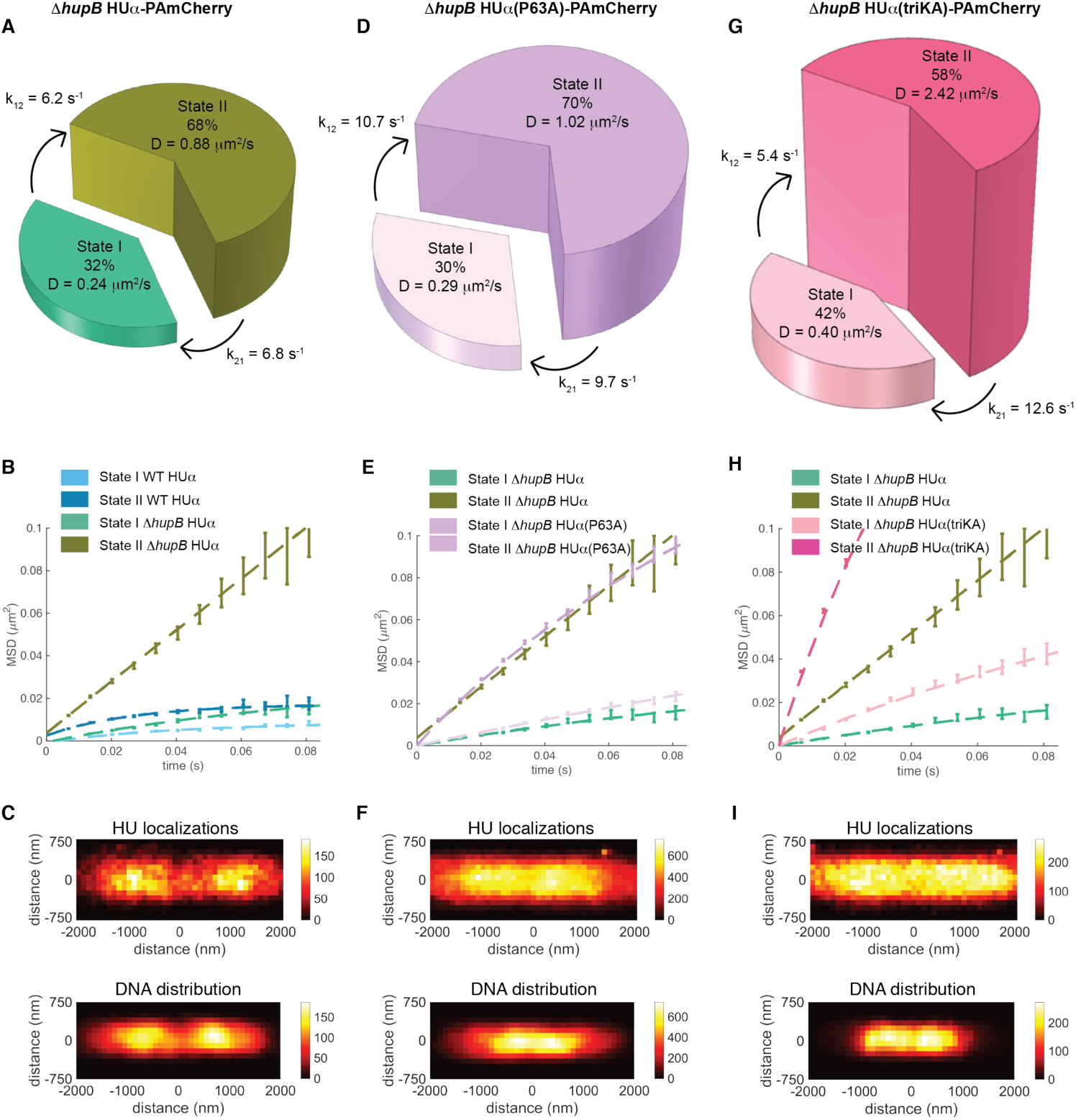
dynamics of HU when HU hetero-dimerization was perturbed demonstrates functional aspects of HU dimers. (A) Two diffusive states of HUα-PAmCherry in Δ*hupB* background with respective population percentages (size of pie piece), transition rates, and diffusion coefficients (height of pie piece) as identified by the HMM. (B) Mean squared displacement (MSD) plots of State I (mint) and State II (dark olive) Δ*hupB* HUα-PAmCherry trajectories as a function of time compared to State I and II molecules in the WT condition (light and dark blue). (C) Two-dimensional (2D) histogram of all cellular Δ*hupB* HUα-PAmCherry localizations from SMT (top) and aggregated nucleoid morphology from SIM (structured illumination microscopy) imaging (bottom). The pixel size of both was 100 × 100 nm. The top color bar indicated the number of localizations used in each bin for Δ*hupB* HUα-PAmCherry (total 30,739 localizations). The bottom color bar indicated the normalized fluorescence level (in arbitrary units) of the nucleoid-intercalating dye Hoechst 33342 (total 20 fluorescence images). (D) Two diffusive states of Δ*hupB* HUα(P63A)-PAmCherry with respective population percentages (size of pie piece), transition rates, and diffusion coefficients (height of pie piece) as identified by the HMM. (E) Mean squared displacement (MSD) plots of State I (light purple) and State II (dark purple) Δ*hupB* HUα(P63A)-PAmCherry trajectories as a function of time compared to State I and II molecules in the WT condition (light and dark blue). (F) Two-dimensional (2D) histogram of all Δ*hupB* HUα(triKA)-PAmCherry localizations from SMT (top) and aggregated nucleoid morphology from SIM (structured illumination microscopy) imaging (bottom). The pixel size of both was 100 × 100 nm. The top color bar indicated the number of localizations used in each bin for Δ*hupB* HUα(P63A)-PAmCherry (total 173,740 localizations). The bottom color bar indicated the normalized fluorescence level (in arbitrary units) of the nucleoid-intercalating dye Hoechst 33342 (total 20 fluorescence images). (G) Two diffusive states of Δ*hupB* HUα(triKA)-PAmCherry with respective population percentages (size of pie piece), transition rates, and diffusion coefficients (height of pie piece) as identified by the HMM. (H) Mean squared displacement (MSD) plots of State I (light pink) and State II (dark pink) Δ*hupB* HUα(triKA)-PAmCherry trajectories as a function of time compared to State I and II molecules in the WT condition (light and dark blue). (I) Two-dimensional (2D) histogram of all Δ*hupB* HUα(triKA)-PAmCherry localizations from SMT (top) and aggregated nucleoid morphology from SIM (structured illumination microscopy) imaging (bottom). The pixel size of both was 100 × 100 nm. The top color bar indicated the number of localizations used in each bin for HUα(triKA)-PAmCherry (total 80,749 localizations). The bottom color bar indicated the normalized fluorescence level (in arbitrary units) of the nucleoid-intercalating dye Hoechst 33342 (total 20 fluorescence images).

Interestingly, the apparent diffusion coefficients and transition kinetics of HUα_2_ observed in Δ*hupB* HUα-PAmCherry cells resembled remarkably those of HUα(P63A)-PAmCherry molecules with (Figure 2D) or without HUβ (Figure 3D, E). The similarities suggest that mutation of P63A in the HUαβ heterodimer likely abolishes the transitory interactions of the heterodimer with chromosomal dsDNA, or causes HUαβ to dissociate so only HUα_2_ homodimer could form. Recalling that HUα(P63A) is implicated in recognizing pre-bent or distorted DNAs in the forms of replication or transcription intermediates, it is possible that a major function of the HUαβ heterodimer is to interact with these specific DNA structures through P63-mediated dynamics, which may play an important role in compacting the nucleoid.

Next, we monitored the dynamics of HUα(triKA)-PAmCherry homodimers in the *ΔhupB* background. We observed the largest increases in the diffusion coefficients (0.4 ± 0.01 μm^2^/s and 2.42 ± 0.03 μm^2^/s, μ ± s.e.m., n = 58,867 displacements from 68 cells) of two significant populations (42% and 58%, Figure 3G). Consistent with the large diffusion coefficients, HUα(triKA)-PAmCherry homodimers were largely cytoplasmic (Figure 3I), mimicking that of non-DNA-interacting protein molecules freely diffusing through the nucleoid. Accompanying this dramatically abolished chromosomal DNA interactions was the more compacted nucleoid of HUα(triKA)_2_ cells compared to that of HUα_2_ homodimer cells, suggesting a role of the dynamic interactions of HU (mediated by triK) with chromosomal DNA in decondensing the nucleoid. This role is also consistent with as we observed on the HUα(triKA)β heterodimer (Figure 2A, Figure S7B). Interestingly, HUα(P63A)_2_ cells also showed more compacted nucleoids compared to HUα_2_ cells, albeit to a less degree than HUα(triKA)_2_ cells (Figure S7B, Table S7). Importantly, both HUα(triKA)_2_ and HUα(P63A)_2_ cells had abnormal, single-lobed nucleoid morphology (Figure 3F, 3I) and severe cell growth defects (Figure S9, S12, Table S4), suggesting that the presence of at least a WT copy of either α or β subunit is important for decondensing the nucleoid, which is required for proper chromosomal DNA segregation.

### HU dynamics are not significantly affected by the loss of naRNAs or nucleoid morphology

HU has been shown to bind many RNA species^13, 28–30^. In particular, a set of highly homologous small non-coding naRNAs expressed from the *REP325* element was shown to bind to HU and subsequently deposited onto cruciformed DNAs to condense relaxed plasmid DNA *in vitro*^30, 31^. Cells deleted of the *REP325* element exhibited expanded nucleoids^30^ (Figure S7), a phenotype which can be reversed by exogenous expression of *naRNA4*^30^. To determine if HU’s binding to naRNAs contribute to its DNA interaction dynamics, we constructed a Δ*REP325* strain in which the entire *REP325* locus was deleted. We then integrated the *hupA-PAmcherry* fusion gene into the Δ*REP325* strain replacing the endogenous *hupA* gene. Single-molecule fluorescence *in-situ* hybridization (smFISH) confirmed that in the Δ*REP325* strain there was a significant reduction of the corresponding naRNAs (Figure S10). However, single-molecule tracking of HUα-PAmCherry in the Δ*REP325* background only showed slight changes (Figure 4A). The two diffusion populations of HUα-PAmCherry had nearly identical apparent diffusion coefficients compared to WT cells; the population percentage of State I molecules and transition kinetics showed changes on the order of 15 – 30% (Figure 3A, Table 1). The MSD plots of both State I and State II HUα-PAmCherry molecules in the *ΔREP325* strain showed relatively larger sub-diffusion zones compared to those of WT cells (Figure 4B, Table S5), consistent with slightly more dispersed cellular distribution of HUα-PAmCherry localizations (Figure 4C). However, the nucleoid volume of the *ΔREP325* was larger than WT cells and comparable to *hupA(P63A)-PAmCherry* cells (Figure S7B). These results indicate that HU’s diffusive dynamics in the nucleoid are not altered by the binding of naRNAs, but naRNAs deposited by HU on the specific structures of chromosomal DNA, likely mediated by P63, facilitate nucleoid condensation, as previous studies suggested^30, 31^.

**Figure 4:**
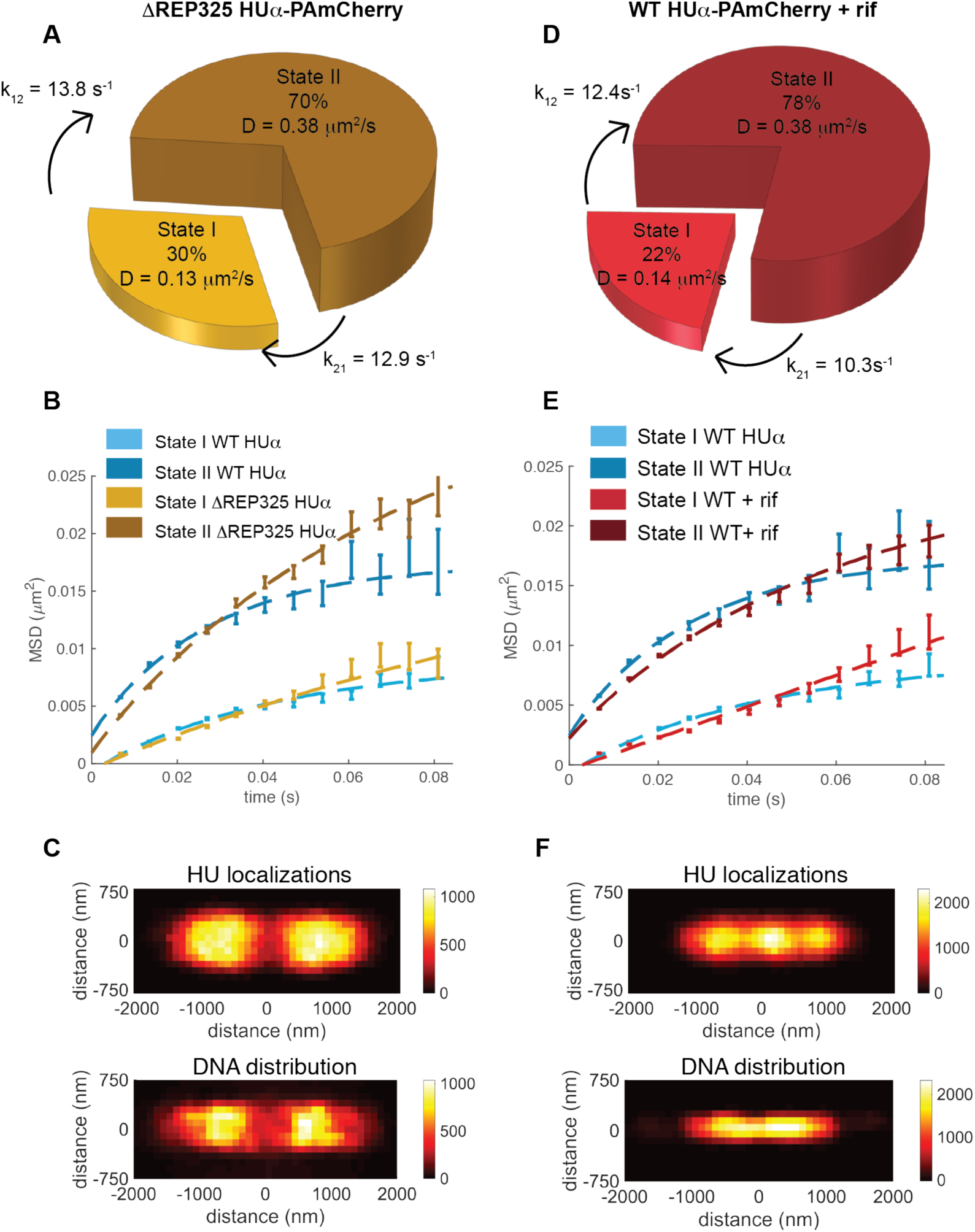
Depletion of ncRNAs did not affect HU dynamics significantly. (A) Two diffusive states of HUα-PAmCherry in Δ*REP325* background with respective population percentages (size of pie piece), transition rates, and diffusion coefficients (height of pie piece) as identified by the HMM. (B) Mean squared displacement (MSD) plots of State I (light gold) and State II (dark gold) Δ*REP325* HUα-PAmCherry trajectories as a function of time compared to State I and II molecules in the WT condition (light and dark blue). (C) Two-dimensional (2D) histogram of all cellular Δ*REP325* HUα-PAmCherry localizations from SMT (top) and aggregated nucleoid morphology from SIM (structured illumination microscopy) imaging (bottom). The pixel size of both was 100 × 100 nm. The top color bar indicated the number of localizations used in each bin for Δ*REP325* HUα-PAmCherry (total 108,846 localizations). The bottom color bar indicated the normalized fluorescence level (in arbitrary units) of the nucleoid-intercalating dye Hoechst 33342 (total 20 fluorescence images). (D) Two diffusive states of HUα-PAmCherry under rifampicin treatment with respective population percentages (size of pie piece), transition rates, and diffusion coefficients (height of pie piece) as identified by the HMM. (E) Mean squared displacement (MSD) plots of State I (light red) and State II (dark red) rifampicin-treated HUα-PAmCherry trajectories as a function of time compared to State I and II molecules in the WT condition (light and dark blue). (F) Two-dimensional (2D) histogram of all cellular rifampicin-treated HUα-PAmCherry localizations from SMT (top) and aggregated nucleoid morphology from SIM (structured illumination microscopy) imaging (bottom). The pixel size of both was 100 × 100 nm. The top color bar indicated the number of localizations used in each bin for HUα-PAmCherry (total 137,019 localizations). The bottom color bar indicated the normalized fluorescence level (in arbitrary units) of the nucleoid-intercalating dye Hoechst 33342 (total 20 fluorescence images).

Next, we reasoned that because there are hundreds of homologous copies of REP elements^51, 52^, and that HU can bind multiple species of naRNAs^29^, it is possible that a single Δ*REP325* background was not sufficient to examine the effect of naRNA binding on HU-DNA interaction dynamics. To test this possibility, we treated HUα-PAmCherry cells with rifampicin (200 µg/mL, 15 min), under which we observed the complete elimination of all cellular naRNAs from REP elements using smFISH (Figure S10). Interestingly, we still observed similar HUα-PAmCherry diffusion dynamics compared to the *ΔREP325* strain (Figure 4D, E, Table 1). These results further confirmed that RNA-binding of HU did not contribute significantly to HU’s interacting dynamics with chromosomal DNA. However, in rifampicin-treated cells, the nucleoid morphology changed significantly (Figure S7); the 2D histogram of the cellular distribution of HUα-PAmCherry localizations exhibited a three-lobed pattern, significantly different from untreated cells (Figure 4F). These results suggested that RNAs could impact nucleoid organization/morphology, but did not affect HU-DNA interaction dynamics. As an additional control for whether altered nucleoid morphology affects HU-DNA interaction dynamics, we treated cells with the translation inhibitor chloramphenicol, which condenses the nucleoid through the inhibition of transcription-coupled translation^53^. We observed very similar diffusion dynamics of HUα-PAmCherry compared to untreated cells, yet highly condensed nucleoid morphology (Figure S11). Correspondingly, the cellular distribution of all HUα-PAmCherry localizations was highly compact as well (Figure S11C). These results demonstrated that the diffusive dynamics of HU and its binding to chromosomal DNAs were not affected by RNA binding or the underlying nucleoid organization.

## Discussion

### HU exhibits transitory binding and unbinding to chromosomal DNAs

Using single molecule tracking of HUα-PAmCherry molecules in live *E. coli* cells, we demonstrated that HUα-PAmCherry had two diffusive states with diffusion coefficients of ∼ 0.1 and ∼ 0.4 µm^2^s^-1^. By comparing HUα-PAmCherry with the diffusion dynamics of another DNA binding protein TetR, we were able to attribute the slow-diffusion population to the DNA-bound state and the fast-diffusion population to the unbound (but nucleoid-constrained) state respectively. An alternative possibility could be that one population was caused by HUα_2_ homodimers and the other by HUαβ heterodimers, which are both present in cells under our growth condition (exponential phase)^7^. This possibility is unlikely though, because in our later experiments when we conducted the same SMT of HUα-PAmCherry in the Δ*hupB* background where only HUα_2_ homodimers could exist, we still observed two distinct diffusive states with larger diffusion coefficients (Figure 3A), suggesting that HUβ must contribute to the dynamics of both slow and fast diffusive states. Mutating three key lysine residues on the surface of HUα important for non-specific binding to linear DNA^32^ effectively abolished the DNA-bound state of HU, implying these residues are critical to HU’s nonspecific DNA-binding activity. Lysine mutant strains with or without HUβ displayed altered cell lengths, slower growth, and nucleoid segregation/cell division defects (Figures S1 and S7, and Table 1), suggesting this binding ability mediated by the triK surface is of critical importance to maintaining healthy cell physiology.

Surprisingly and importantly, we found that HU is highly dynamic and switches rapidly between the two diffusive states. This dynamic behavior of HU is in stark contrast to other DNA-binding proteins, such as histones in eukaryotic cells, or the bacterial NAP H-NS, which exhibit FRAP recoveries on the order of hours^54^ and minutes^55^ respectively, but is consistent with prior ChIP studies of HU in *E. coli*^12^ and the recent findings of HU dynamics in *D. radiodurans*^56^. Historically, HU-mediated nucleoid organization has been proposed to be facilitated by stable or semi-stable nucleosome-like structures, supported by previous *in vitro* work^32, 46, 47^. If these models were indeed relevant *in vivo*, our observations could suggest that HU existing in these multimeric structures are constantly exchanging with the fast diffusing population to allow for rapid reorganization of the local chromosome structure. Regardless, the direct observation of highly dynamic HU proteins demonstrates that HU is distinctively not histone-like and does not play a static, architectural role in chromosome organization.

To explain the seemingly disparate observations of the transitory dynamics of HU with its critical importance to nucleoid organization, we propose a model in which HU, through the collective sum of a large number of weak, transitory interactions with chromosomal DNAs, maintains the viscoelastic properties and fluidity of the bacterial nucleoid (Figure 5). We use these lines of evidence to support our hypothesis: (1) nucleoids of *E. coli* can undergo global changes in morphology in as little as 5s^57^, which presumably necessitates dynamic, transiently bound NAPs and increased DNA flexibility; (2) loss of HU disrupts small DNA loop formation *in vivo*^27^, very likely due to an increase in DNA persistence length as demonstrated by *in vitro* single molecule experiments^23, 58^; (3) HU does not play a static architectural role in chromosome organization, as significant modulation of nucleoid condensation through drug treatments did not significantly alter HU dynamics or localization (Figure 3 and S11), unlike the loss of chromosomal dsDNA binding in the triKA mutant (Figure 2); and (4) in the event of HU deletion, our theory predicts a global change in the mechanical properties of the DNA, consistent with the previously observed changes in DNA replication^10^, recombination^17, 59^, gene expression^12^, and nucleoid segregation^60^ in a HU deletion background. The viscoelastic maintenance model has been demonstrated for other systems-for example, the non-specific, transitory interactions of the muscle protein α-actinin modulates the mechanical properties of the actin filament network to form a viscoelastic gel^61^. Interestingly, actin filament networks, due to their dynamic nature, demonstrate properties of liquid phase separation^62^. In the case of HU, this could suggest that the dynamic nature of HU binding to chromosomal DNAs could maintain the nucleoid as a separate “macro-compartment” that is physically distinct from the cytosol, and could explain why HU deletion strains have increased nucleoid volumes. Overall, our findings underscore the importance of weak, non-specific binding on the biophysical properties of the cell and provide strong evidence for the contribution of weak binding to the maintenance of the mechanical properties of the chromosome.

**Figure 5:**
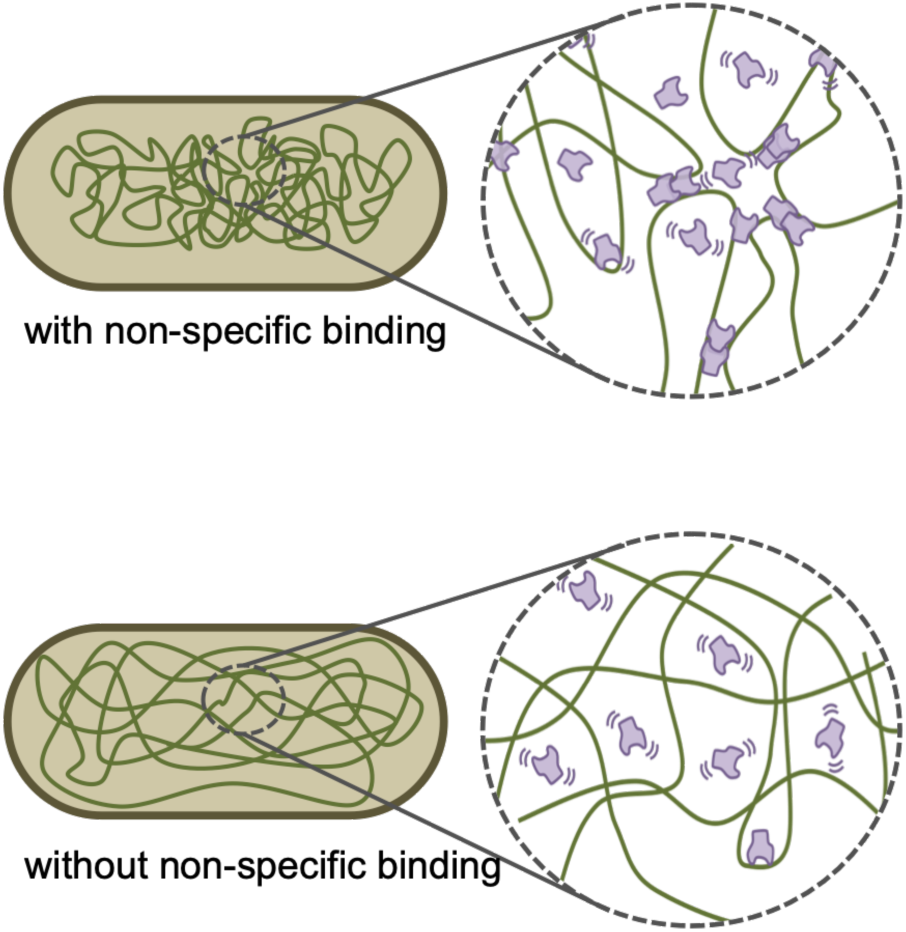
transient, nonspecific interactions of HU with chromosomal DNAs maintains the viscoelastic properties of the nucleoid. We unambiguously identified the transient, dynamic nature of HU with chromosomal DNAs. This observation in combination with HU’s role in global topological processes such as replication, recombination, or nucleoid segregation, suggest a model whereby the transient, nonspecific HU-DNA interactions contribute a collective “force” that alters the physical properties of the nucleoid to facilitate nucleoid formation, distant DNA-DNA contacts, and increased DNA flexibility for the aforementioned global topological processes.

### HUα_2_ dimers have weaker apparent affinity for chromosomal DNAs than HUαβ heterodimers

Because *E. coli* express HUα_2_ and HUαβ at different levels during cell growth, it was thought these differential forms must exert differential effects on nucleoid organization. When we imaged HUα-PAmCherry molecules in a *ΔhupB* background (i.e. exclusively HUα_2_ dimers), we observed increased diffusion coefficients and larger sub-diffusive domains compared to the WT strain, suggesting that HUα_2_ dimers interact with chromosomal dsDNA less frequently. This suggestion is consistent with prior observations that the nucleoid occupies a large volume in cells during the lag and early exponential phases, and then becomes increasingly condensed as cells go into late exponential and stationary phase growth^32^. Additionally, prior biochemical characterization of the various HU dimers has suggested that HUα_2_ has a slightly worse affinity for bulk dsDNA than the HUαβ heterodimer^16^, consistent with our findings. These observations suggest that these relatively small differences in the dynamic interactions of HU dimers can cause global changes in the organization state of the nucleoid.

### Contributions of specific DNA structure-binding of HUα to its diffusive dynamics and nucleoid organization

In contrast to the triple lysine mutant, when we mutated the conserved proline residue important for structure-specific binding such as in homologous recombination intermediates^15^, HU bundling^32^, and transcriptional repression loops^25, 48^, we saw a less dramatic change in HU dynamics. Interestingly, HUα(P63A)-PAmCherry (in the forms of HUα(P63A)β or HUα(P63A)_2_ dimers) in the presence or absence of HUβ displayed surprisingly similar dynamic behavior to HUα_2_ homodimers, suggesting that P63A might have disrupted HUαβ heterodimer formation or function. In addition, HUα(P63A)-PAmCherry with HUβ had normal cell lengths and only a minor increase in cell doubling time, while HUα(P63A)-PAmCherry in the Δ*hupB* background had extremely elongated cell lengths and longer doubling times. However, cells regained healthy physiology when grown at 37°C (Figure S9) suggesting the P63 residue may be important for programmed gene expression responses such as cold shock, as previously implicated^63^. Interestingly, we observed that in the Δ*hupB hupA(triKA)-PAmCherry* strain, the nucleoid was significantly more condensed than the Δ*hupA* Δ*hupB* strain, implying that the P63 residues are capable of condensing the nucleoid through a mechanism that may be dependent on specific DNA structure-binding mediated by the lysine-rich surface.

### RNA binding and nucleoid structure had limited impact HU dynamics

HU is known to bind to several species of RNAs, particularly the *REP325* naRNAs^29–31^. Our observations of HUα-PAmCherry both in a Δ*REP325* strain and under total RNA depletion via rifampicin treatment showed a moderately decreased population of slow, DNA-bound HU and a corresponding change in its DNA association rate. Curiously, we found that in the Δ*REP325* strain, HU remained nucleoid-localized but had slightly increased nucleoid volume. Furthermore, in rifampicin treatment, nucleoids were highly modulated (Figure S7), yet HU remained nucleoid-localized and its dynamics were nearly identical to HU in the Δ*REP325* strain. These results are surprising given previous reports that loss of naRNAs results in loss of specific long-distance DNA contacts^31^, but suggests RNA binding to HU does not influence HU dynamics. Furthermore, HU dynamics were not significantly affected by changing the underlying nucleoid structure, such as with rifampicin or chloramphenicol treatment, suggesting that HU may not be responsible for maintaining a stable nucleoid architecture, but rather on maintaining a dynamic chromosome by influencing the viscoelastic properties of the nucleoid as we proposed in the model.

## Materials and Methods

### Cell growth

Strains were inoculated from single colonies from freshly-streaked LB plates into EZ Rich Defined Media (EZRDM, Teknova) using 0.4% glucose as the carbon source with chloramphenicol antibiotic (Sigma-Aldrich C0378) added to 150 µg/mL, and carbinicillen antibiotic (Sigma-Aldrich C3416) added to 60 µg/mL respectively. Cells were grown overnight at room temperature (RT), shaking at 240 rpm. The following morning, saturated cultures were re-inoculated into fresh EZRDM and grow for several hours at RT until mid-log phase growth (OD600 ∼0.4). For rifampicin drug treatment, rifampicin antibiotic (Sigma-Aldrich R3501) was added to a final concentration of 200 µg/mL for 15 minutes before cells were harvested for imaging (see Materials and Methods: SMT data collection below). For chloramphenicol drug treatment, additional chloramphenicol was added to a final concentration of 600 µg/mL for 30 minutes before cells were harvested for imaging. For both drug treatments, the drug was added when cells were at mid-log phase (OD600 ∼ 0.4). For imaging of the *galP* fluorescent reporter operator system, cells were spun down at 4.5 rcf for 5 minutes and resuspended in EZRDM media with 0.4% arabinose to induce expression of the reporter gene. Induction was done at RT for 2 hours with shaking, then cells were again spun down at 4.5 rcf for 5 minutes, washed twice with fresh EZRDM media, then resuspended in fresh EZRDM media and grown at 30°C for 1 hour with shaking to allow for maturation of the mCherry fluorophore, after which cells were harvested as described in the section below.

### Growth curves

For growth curves, distinct colonies on agar plates were inoculated into 2mL of EZRDM media for each strain and grown overnight at 24°C, shaking at 240 rpm. The following afternoon, saturated cultures were diluted 1 to 200 into a 96 well plate (100uL volume total for each well) along with a few wells with EZRDM media alone to serve as a control. The 96 well plate was placed into a microplate reader (Life Sciences Tecan) and the absorbance at 600nm was measured every 30 minutes for a total of 12 hours. Reads were normalized against the control wells.

### Construction of strains

Strains used in this study are mentioned in Supplementary Table 1.

The construction of the *HupA-PAmCherry* strain (KB0026) has been described elsewhere^33^. To test the functionality of the PAmCherry tagged HUα in the absence of HUβ, the *ΔhupB* mutation from the donor strain SCV152 was transduced into the *HupA-PAmCherry* strain using standard P1vir transduction protocol. Before P1vir transduction, the chloramphenicol resistance cassette linked to PAmCherry was removed using the temperature sensitive plasmid pCP20^64^ that expresses the recombinase flippase. For this, the *HupA-PAmCherry* strain was transformed with pCP20 plasmid and grown overnight at 42°C to induce kickout of the chloramphenicol resistance cassette flanked by short flippase recognition target (FRT) sites by the recombinase flippase. *hupA(triKA)-PAmCherry* (SCV149) *and hupA(P63A)-PAmCherry* (SCV146) were constructed by lambda red recombineering using pSIM6 plasmid^65^. In the first recombineering step, a kan-pBAD-ccdB cassette was introduced immediately after the stop codon of wild-type *hupA* gene of MG1655. In the second recombineering step, the wild-type *hupA* gene was replaced with the *hupA* gene encoding HUα(P63A) or HUα*(triKA)* protein. A synthetic double stranded DNA synthesized by Integrated Technologies (IDT) with the mutation was used for the recombination. The presence of the mutation was confirmed by Sanger sequencing. In the third recombineering step, PA-mCherry linked with chloramphenicol resistance cassette was amplified from KB0026 and was fused in-frame to the *hupAP63A* gene after the last codon. The Δ*hupB*726::Kan^R^ from the strain JW0430-3, which was obtained from Coli Genetic Stock Center (CGSC), Yale University, was transduced into SCV146 and SCV149 to yield the strains Δ*hupB hupA(P63A)-PAmCherry* (SCV150) and Δ*hupB hupA(triKA)-PAmCherry (SCV153). For the HupA-PAmCherry* strain in the MG1655 background (SCV148), PAmCherry linked with chloramphenicol resistance cassette was fused in-frame to the wild-type *hupA* gene of MG1655 after the last codon. Subsequently, the Δ*hupB*726::Kan^R^ from the strain JW0430-3 was transduced into SCV148 to yield the strain Δ*hupB hupA-PamCherry* (SCV152).

### Mu phage assay

Spot dilution plates were performed initially to estimate proper Mu phage lysate dilution to use for phage plaque counting. Cells were grown overnight in Luria Broth (LB) at 37°C, and the next day reinoculated into LB with 1 mM CaCl_2_ and 2.5 mM MgCl_2_. Cells were grown until O.D. ∼ 0.4, then spun down and resuspended so the final O.D. for later use in the protocol is ∼ 1. Next, 100 µl of cells were mixed with 10 µl of the appropriate dilution of Mu phage lysate. This was mixed quickly into 3 ml of 0.7% top agar (warmed to 55 °C) and immediately poured onto TB plates (warmed to 37°C) and redistributed evenly prior to hardening. Plates were let dry at RT and incubated overnight at 37°C. The next day, plates were examined and the plaques were counted by eye. The PFU/ml (plaque forming units, Mu/mL) was calculated using the following equation:

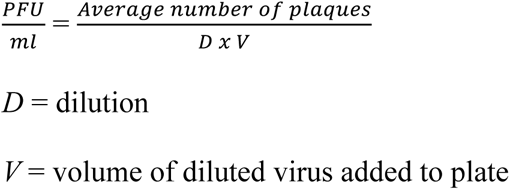

### Mini-P1 plasmid transformation assay

Electrocompetent cells were made using standard protocol. In brief, 6 ml of *E. coli* cells were grown until O.D. ∼0.4 in LB, the cultures were put on ice for 30 min. Cells were spun down at 4100 rpm at 4°C for 10 min, cell pellet was resuspended in 6 ml ice cold H_2_O. Cells were spun down again and resuspended in 3 ml of ice cold 10% glycerol. This was repeated once more, then cells were resuspended in 1.5 ml ice cold 10% glycerol and spun down again. The final cell pellet was resuspended in ∼300 µl of GYT media, and 6 individual aliquots of 50 µl of competent cells were made. Cells were quickly frozen in dry ice, and stored at −80°C until later use.

The same batch of competent cells were used for all the transformation experiments for consistency. The pUC19 transformation control was performed in order to normalize the transformation efficiency of competent cells between the different *E. coli* strains. An appropriate volume of transformation outgrowth was plated on appropriate antibiotic selection LB agar plates and incubated O/N at 37°C. The next day, the numbers of colonies on agar plates were counted and recorded. The mini-P1 CFU/µg DNA was normalized to the same strain’s pUC19 CFU/µg DNA to account for competent cell transformation efficiency differences. The WT MG1655 strain was set as the reference standard for comparison with other *E. coli* strains.

### SMT data collection

Agarose gel pads were made using a low-melting-temperature agarose (SeaPlaque, Lonza) and EZRDM media, with agarose concentration at 3% w/v. 1.5mL of cells were harvested by spinning down in a bench-top microcentrifuge at 4.5 rcf for 5 minutes and resuspended in ∼100µL of fresh EZRDM. Cells were pipetted onto agarose pads and sandwiched between the agarose pad and a #1 coverslip. Immobilized cells were imaged on an Olympus IX71 inverted microscope with a 100x oil objective (PN, NA =1.45) with 1.6x additional magnification. Photons from cells were collected with an Andor EMCCD camera using MetaMorph imaging software (Molecular Devices). Fluorescence from cells was obtained using solid-state lasers at 405nm and 568nm wavelengths (Coherent). All SMT images were collected using 5ms exposure with 1.74ms of cycle time for a total frame length of 6.74ms. Three movies consisting of 2500 frames each were taken for each cell. No cell was imaged longer than five minutes to avoid phototoxicity effects. Activation of the fluorescent proteins was continuous throughout imaging, and no changes were made to either the activation or excitation power throughout the imaging session.

### SMT analysis

For single molecule tracking analysis, tiff stacks of cell images were imported into the single molecule tracking software UTrack version 3.1^66^ within the Matlab 2017a software. We performed the detection and tracking of molecules on individual movies using the Gaussian Mixture Model setting within the UTrack software. For detection, an α value of 0.01 was used; for frame-to-frame linking, 15 frames was the maximum time and 0 to 2 pixels (or up to 6 pixels with increased linking cost) was the maximum distance to link trajectories together. Linked trajectories were then filtered by their intensity (between 0.5-1.5x the average single molecule intensity to exclude molecules that had temporarily colocalized) and their localization within cells (i.e. any trajectory outside of cells was excluded) and exported as MATLAB *mat* files. Trajectories were analyzed using custom in-house code (available upon request) to assess their diffusive properties. Exact equations used to analyze the trajectories are explained in more detail in the supplemental information. To build the HMM, we installed and used the software vbSPT, version 1.1 on MATLAB version 2014a^34^. Cells were rotated such that the long cell-axis corresponded to the x-axis. We used only consecutive frame trajectories and only displacements in the x-direction for analysis to limit the effects of confinement due to the small size of the bacterial cell.

### Localizations of HU-PAmCherry

Detections of HU-PAmCherry were determined as above using the UTrack software. Cells were rotated such that the long cell axis corresponded to the x-axis, and the middle of the cell corresponded to the origin. We determined the spatial coordinates based on this axis and added spatial coordinates of all cells together. For single molecule tracking experiments, cells were chosen to fit within the imaging view (40 × 40 pixels) and thus have a narrow distribution of cell lengths and widths. We binned these displacements into 100 × 100 nanometer bins and applied the red-hot colormap for viewing. Images were cropped to 4 microns long and 1.5 microns wide to enable direct comparison across conditions.

### Imaging of cell nucleoids

To image nucleoids, strains were grown as above, then stained with 5 µM of Hoechst 33342 dye for 10 minutes. Cells were washed twice with fresh media to remove unincorporated dye, then fixed in 3.7% paraformaldehyde (15 min). Cells were washed twice with PBS and resuspended in a final volume of 200uL and stored in a light-proof box at 4°C until ready to image. Cells were kept in the 4°C box for no longer than one week. Fixed and stained cells were mixed 1:1 with anti-fading solution (20% NPG (n-propyl gallate), 60% glycerol, 1x PBS) and adhered to coverslips treated with 0.01% poly-L-lysine solution. Coverslips were sealed onto a microscope slide using clear nail polish and imaged on the GE OMX SR structured illumination microscope (excitation: 405, camera channel: 488, exposure 50ms, 5% of highest laser intensity). Images were reconstructed using the standard parameters for the GE OMX SR microscope within the GE SRx software.

### Analysis of nucleoid images

For each strain, individual cell images were cropped and rotated such that the long axis of the cell corresponded to the x-axis. Black space was manually added to the edges of images to center cells, such that the middle of the cell corresponded to the axis origin. Each set of images for each strain was then added together and re-binned such that each bin corresponded to 100 × 100 nm to allow for direct comparison to HUα-PAmCherry localizations. All images were cropped to 4 microns long and 1.5 microns wide. For better visual comparison, matrix values were normalized to 1 and then multiplied by the total localizations of HUα-PAmCherry.

To determine the area of the nucleoid, we first normalized images using HUα-PAmCherry as a template using histogram matching combined with gradient information^67^ to account for intensity and noise differences inherent from day-to-day imaging. Next, we determined a threshold based on Otsu’s method (graythresh.m in MATLAB), ensuring the effectiveness was greater than 0.85, to generate a binary image. The nucleoid area was then the number of pixels above the calculated threshold times the pixel size (40 × 40 nm). To estimate the error, we used bootstrapping (boostrap.m in MATLAB) to re-sample the average nucleoid intensities and repeated the area measurement as before. Bootstrapping was completed 100 times per strain. These bootstrapped means were used to perform a one-way ANOVA test to determine if these nucleoid areas were statistically significant from each other.

### Total cellular RNA extraction

Cell growth conditions are same as described above. Starting with an identical number of cells for WT condition (no drug treatment), rifampicin treatment (200 µg/mL, 60 min) and chloramphenicol treatment (600 µg/mL, 60 min). Take volume of cells to an O.D. equivalent of 8.5, pellet cells (10 min, 4100 rpm, 4°C), resuspend the pellet in 1 mL 1xPBS, and transfer to an Eppendorf tube. The cells were pelleted again (2 min, 14000 rpm, 4°C), the pellet was quickly frozen on dry ice and kept at −80°C until the later steps. For RNA extraction, 600 µL of Trizol was added to the cell pellet after thawing, then the protocol from the Direct-zol RNA Miniprep Kit (Zymo Research) was followed, an in-column DNase I digestion was performed as suggested for 15 min. Finally, RNA was eluted with 50 µl of DNase/RNase free H_2_O and quantified by measuring on Nanodrop.

### RNA FISH

Cells were grown the same as for imaging (described above; grown in rich defined media at 24°C). At mid-log phase growth corresponding to OD_600_ ∼ 0.4, cells were fixed in 2.6% paraformaldehyde, 0.8% glutaraldehyde in 1 × PBS for 15 minutes at room temperature. Cells were then washed twice in 1 × PBS and resuspended into a small volume of GTE buffer (2mL of cells were spun into 50uL final volume). Coverslips were treated with 0.1% poly-L-lysine for 5 to 10 minutes, then rinsed and dried. Cells were spotted onto treated coverslips and left to dry for 20 minutes. Non-adhered cells were washed away with PBS. Cells were then treated with ice-cold 70% ethanol and placed at −20°C for 10 minutes to permeabilize the cells. After permeabilization, cells were washed twice in 2 × SSCT (2 × SSC + 0.1% Tween-20) for five minutes each wash. Cells were then pre-hybridized in pre-hybridization solution (2 × SSC, 40% formamide, 0.1% Tween-20) for 30 to 45 minutes at 37°C. Pre-hybridization solution was removed, then a small volume of hybridization solution (2 × SSC, 40% formamide, dextran sulfate, yeast tRNAs) with 1µM of Cy3-labelled probe was spotted onto the coverslip, and covered with a clean piece of saran wrap to set and prevent evaporation. Coverslips were placed in a sterile petri dish and submerged in a water bath set to 94°C for two minutes, then immediately placed at 42°C for 15 to 18 hours to hybridize. The probe sequence used was: Cy3-GTTGCCGGATGCGGCGTAAACGCCTTATCCGGCC. Cells were washed in 40% wash solution (2 × SSC, 40% formamide, 0.1% Tween-20) for 30 minutes at 37°C, then washed in 20% wash solution (2 × SSC, 20% formamide, 0.1% Tween-20) for 10 minutes at room temperature, then washed in 2 × SSCT solution for 10 minutes at room temperature, and finally transferred to 2 × SSC solution until imaging. Coverslips were mounted onto imaging chambers and anti-fading media was perfused into the system. Cells were imaged on an Olympus IX71 inverted microscope with a 100x oil objective (NA =1.45) with 1.6x additional magnification. Photons from cells were collected with an Andor EMCCD camera using MetaMorph imaging software (Molecular Devices). Fluorescence from cells was obtained using solid-state lasers at 568nm wavelength (Coherent). Z-stacks of cells were collected using 50ms exposure per frame with an EM gain of 300. Z-stacks were imported to ImageJ, where cells were manually identified. Fluorescence in each cell was corrected by subtraction of the background (a random area devoid of cells for each image). Although the sequence was against *REP325 naRNA4*, signal remained above background in the Δ*REP325* background strain due to high homology of the *REP325* RNAs with hundreds of other *REP* elements in the cell. For each of the two biological replicates, over 100 cells were analyzed.

## References

1. Sinden, R. R. & Pettijohn, D. E. Chromosomes in living Escherichia coli cells are segregated into domains of supercoiling. Proc Natl Acad Sci USA 78, 224–228 (1981).

2. Boccard, F., Esnault, E. & Valens, M. Spatial arrangement and macrodomain organization of bacterial chromosomes. Mol. Microbiol. 57, 9–16 (2005).

3. Browning, D. F., Grainger, D. C. & Busby, S. J. Effects of nucleoid-associated proteins on bacterial chromosome structure and gene expression. Current Opinion in Microbiology 13, 773–780 (2010).

4. Badrinarayanan, A., Le, T. B. K. & Laub, M. T. Bacterial chromosome organization and segregation. Annu. Rev. Cell Dev. Biol. 31, 171–199 (2015).

5. Valens, M., Penaud, S., Rossignol, M., Cornet, F. & Boccard, F. Macrodomain organization of the Escherichia coli chromosome. The EMBO Journal 23, 4330–4341 (2004).

6. Ali Azam, T., Iwata, A., Nishimura, A., Ueda, S. & Ishihama, A. Growth phase-dependent variation in protein composition of the Escherichia coli nucleoid. Journal of Bacteriology 181, 6361–6370 (1999).

7. Claret, L. & Rouviere-Yaniv, J. Variation in HU composition during growth of Escherichia coli: the heterodimer is required for long term survival. J. Mol. Biol. 273, 93–104 (1997).

8. Broyles, S. S. & Pettijohn, D. E. Interaction of the Escherichia coli HU protein with DNA. Evidence for formation of nucleosome-like structures with altered DNA helical pitch. J. Mol. Biol. 187, 47–60 (1986).

9. Rouviere-Yaniv, J., Yaniv, M. & Germond, J. E. E. coli DNA binding protein HU forms nucleosomelike structure with circular double-stranded DNA. Cell 17, 265–274 (1979).

10. Dixon, N. E. & Kornberg, A. Protein HU in the enzymatic replication of the chromosomal origin of Escherichia coli. Proc Natl Acad Sci USA 81, 424–428 (1984).

11. Bonnefoy, E. & Rouviere-Yaniv, J. HU, the major histone-like protein of E. coli, modulates the binding of IHF to oriC. The EMBO Journal 11, 4489–4496 (1992).

12. Prieto, A. I. et al. Genomic analysis of DNA binding and gene regulation by homologous nucleoid-associated proteins IHF and HU in Escherichia coli K12. 40, 3524–3537 (2012).

13. Balandina, A., Claret, L., Hengge-Aronis, R. & Rouviere-Yaniv, J. The Escherichia coli histone-like protein HU regulates rpoS translation. Mol. Microbiol. 39, 1069–1079 (2001).

14. Kar, S., Edgar, R. & Adhya, S. Nucleoid remodeling by an altered HU protein: reorganization of the transcription program. Proc Natl Acad Sci USA 102, 16397–16402 (2005).

15. Kamashev, D. & Rouviere-Yaniv, J. The histone-like protein HU binds specifically to DNA recombination and repair intermediates. The EMBO Journal 19, 6527–6535 (2000).

16. Pinson, V., Takahashi, M. & Rouviere-Yaniv, J. Differential binding of the Escherichia coli HU, homodimeric forms and heterodimeric form to linear, gapped and cruciform DNA. J. Mol. Biol. 287, 485–497 (1999).

17. Li, S. & Waters, R. Escherichia coli strains lacking protein HU are UV sensitive due to a role for HU in homologous recombination. Journal of Bacteriology 180, 3750–3756 (1998).

18. Malik, M., Bensaid, A., Rouviere-Yaniv, J. & Drlica, K. Histone-like protein HU and bacterial DNA topology: suppression of an HU deficiency by gyrase mutations. J. Mol. Biol. 256, 66–76 (1996).

19. Wada, M., Kano, Y., Ogawa, T., Okazaki, T. & Imamoto, F. Construction and characterization of the deletion mutant of hupA and hupB genes in Escherichia coli. J. Mol. Biol. 204, 581–591 (1988).

20. Huisman, O. et al. Multiple defects in Escherichia coli mutants lacking HU protein. Journal of Bacteriology 171, 3704–3712 (1989).

21. Bensaid, A., Almeida, A., Drlica, K. & Rouviere-Yaniv, J. Cross-talk between topoisomerase I and HU in Escherichia coli. J. Mol. Biol. 256, 292–300 (1996).

22. Lal, A. et al. Genome scale patterns of supercoiling in a bacterial chromosome. Nat Commun 7, 11055 (2016).

23. van Noort, J., Verbrugge, S., Goosen, N., Dekker, C. & Dame, R. T. Dual architectural roles of HU: formation of flexible hinges and rigid filaments. Proc Natl Acad Sci USA 101, 6969– 6974 (2004).

24. Becker, N. A., Kahn, J. D. & Maher, L. J. Effects of nucleoid proteins on DNA repression loop formation in Escherichia coli. 35, 3988–4000 (2007).

25. Aki, T., Choy, H. E. & Adhya, S. Histone-like protein HU as a specific transcriptional regulator: co-factor role in repression of gal transcription by GAL repressor. Genes to Cells 1, 179–188 (1996).

26. Zimmerman, J. & Maher, L. J. Transient HMGB protein interactions with B-DNA duplexes and complexes. Biochem. Biophys. Res. Commun. 371, 79–84 (2008).

27. Becker, N. A., Kahn, J. D. & Maher, L. J. Bacterial repression loops require enhanced DNA flexibility. J. Mol. Biol. 349, 716–730 (2005).

28. Balandina, A., Kamashev, D. & Rouviere-Yaniv, J. The bacterial histone-like protein HU specifically recognizes similar structures in all nucleic acids. DNA, RNA, and their hybrids. J. Biol. Chem. 277, 27622–27628 (2002).

29. Macvanin, M. et al. Noncoding RNAs binding to the nucleoid protein HU in Escherichia coli. Journal of Bacteriology 194, 6046–6055 (2012).

30. Qian, Z. et al. A New Noncoding RNA Arranges Bacterial Chromosome Organization. MBio 6, e00998–15 (2015).

31. Qian, Z., Zhurkin, V. B. & Adhya, S. DNA-RNA interactions are critical for chromosome condensation in Escherichia coli. Proc. Natl. Acad. Sci. U.S.A. 114, 12225–12230 (2017).

32. Hammel, M. et al. HU multimerization shift controls nucleoid compaction. Sci Adv 2, e1600650 (2016).

33. Wang, S., Moffitt, J. R., Dempsey, G. T., Xie, X. S. & Zhuang, X. Characterization and development of photoactivatable fluorescent proteins for single-molecule-based superresolution imaging. Proc. Natl. Acad. Sci. U.S.A. 111, 8452–8457 (2014).

34. Persson, F., Lindén, M., Unoson, C. & Elf, J. Extracting intracellular diffusive states and transition rates from single-molecule tracking data. Nat Meth 10, 265–269 (2013).

35. Vrljic, M., Nishimura, S. Y. & Moerner, W. E. Single-molecule tracking. Methods Mol. Biol. 398, 193–219 (2007).

36. Jen-Jacobson, L., Engler, L. E. & Jacobson, L. A. Structural and thermodynamic strategies for site-specific DNA binding proteins. Structure 8, 1015–1023 (2000).

37. Lau, I. F. et al. Spatial and temporal organization of replicating Escherichia coli chromosomes. Mol. Microbiol. 49, 731–743 (2003).

38. Hensel, Z., Weng, X., Lagda, A. C. & Xiao, J. Transcription-factor-mediated DNA looping probed by high-resolution, single-molecule imaging in live E. coli cells. PLoS Biol. 11, e1001591 (2013).

39. Javer, A. et al. Short-time movement of E. coli chromosomal loci depends on coordinate and subcellular localization. Nat Commun 4, 3003 (2013).

40. Elowitz, M. B., Surette, M. G., Wolf, P. E., Stock, J. B. & Leibler, S. Protein mobility in the cytoplasm of Escherichia coli. Journal of Bacteriology 181, 197–203 (1999).

41. Elf, J., Li, G.-W. & Xie, X. S. Probing transcription factor dynamics at the single-molecule level in a living cell. Science 316, 1191–1194 (2007).

42. Stracy, M. et al. Live-cell superresolution microscopy reveals the organization of RNA polymerase in the bacterial nucleoid. Proc. Natl. Acad. Sci. U.S.A. 112, E4390–9 (2015).

43. Stracy, M. et al. Single-molecule imaging of UvrA and UvrB recruitment to DNA lesions in living Escherichia coli. Nat Commun 7, 12568 (2016).

44. Fei, J., et al. RNA biochemistry. Determination of in vivo target search kinetics of regulatory noncoding RNA. Science 347, 1371–1374 (2015).

45. Dayton, C. J., Prosen, D. E., Parker, K. L. & Cech, C. L. Kinetic measurements of Escherichia coli RNA polymerase association with bacteriophage T7 early promoters. J. Biol. Chem. 259, 1616–1621 (1984).

46. Sarkar, T., Vitoc, I., Mukerji, I. & Hud, N. V. Bacterial protein HU dictates the morphology of DNA condensates produced by crowding agents and polyamines. 35, 951–961 (2007).

47. Dame, R. T., Hall, M. A. & Wang, M. D. Single-molecule unzipping force analysis of HU-DNA complexes. Chembiochem 14, 1954–1957 (2013).

48. Aki, T. & Adhya, S. Repressor induced site-specific binding of HU for transcriptional regulation. The EMBO Journal 16, 3666–3674 (1997).

49. Swinger, K. K., Lemberg, K. M., Zhang, Y. & Rice, P. A. Flexible DNA bending in HU-DNA cocrystal structures. The EMBO Journal 22, 3749–3760 (2003).

50. Guo, F. & Adhya, S. Spiral structure of Escherichia coli HUalphabeta provides foundation for DNA supercoiling. Proc Natl Acad Sci USA 104, 4309–4314 (2007).

51. Dimri, G. P., Rudd, K. E., Morgan, M. K., Bayat, H. & Ames, G. F. Physical mapping of repetitive extragenic palindromic sequences in Escherichia coli and phylogenetic distribution among Escherichia coli strains and other enteric bacteria. Journal of Bacteriology 174, 4583–4593 (1992).

52. Blattner, F. R. et al. The complete genome sequence of Escherichia coli K-12. Science 277, 1453–1462 (1997).

53. Cabrera, J. E., Cagliero, C., Quan, S., Squires, C. L. & Jin, D. J. Active transcription of rRNA operons condenses the nucleoid in Escherichia coli: examining the effect of transcription on nucleoid structure in the absence of transertion. Journal of Bacteriology 191, 4180– 4185 (2009).

54. Kimura, H. Histone dynamics in living cells revealed by photobleaching. DNA Repair (Amst.) 4, 939–950 (2005).

55. Kumar, M., Mommer, M. S. & Sourjik, V. Mobility of cytoplasmic, membrane, and DNA-binding proteins in Escherichia coli. Biophysical Journal 98, 552–559 (2010).

56. Floc’h, K. et al. Cell morphology and nucleoid dynamics in dividing Deinococcus radiodurans. Nat Commun 10, 3815–13 (2019).

57. Fisher, J. K. et al. Four-dimensional imaging of E. coli nucleoid organization and dynamics in living cells. Cell 153, 882–895 (2013).

58. Kundukad, B., Cong, P., van der Maarel, J. R. C. & Doyle, P. S. Time-dependent bending rigidity and helical twist of DNA by rearrangement of bound HU protein. Nucleic Acids Res. 41, 8280–8288 (2013).

59. Dri, A. M., Moreau, P. L. & Rouviere-Yaniv, J. Role of the histone-like proteins OsmZ and HU in homologous recombination. Gene 120, 11–16 (1992).

60. Jaffé, A., Vinella, D. & D’Ari, R. The Escherichia coli histone-like protein HU affects DNA initiation, chromosome partitioning via MukB, and cell division via MinCDE. Journal of Bacteriology 179, 3494–3499 (1997).

61. Xu, J., Wirtz, D. & Pollard, T. D. Dynamic cross-linking by alpha-actinin determines the mechanical properties of actin filament networks. J. Biol. Chem. 273, 9570–9576 (1998).

62. Weirich, K. L. et al. Liquid behavior of cross-linked actin bundles. Proc. Natl. Acad. Sci. U.S.A. 114, 2131–2136 (2017).

63. Giangrossi, M., Giuliodori, A. M., Gualerzi, C. O. & Pon, C. L. Selective expression of the beta-subunit of nucleoid-associated protein HU during cold shock in Escherichia coli. Mol. Microbiol. 44, 205–216 (2002).

64. Cherepanov, P. P. & Wackernagel, W. Gene disruption in Escherichia coli: TcR and KmR cassettes with the option of Flp-catalyzed excision of the antibiotic-resistance determinant. Gene 158, 9–14 (1995).

65. Datta, S., Costantino, N. & Court, D. L. A set of recombineering plasmids for gram-negative bacteria. Gene 379, 109–115 (2006).

66. Jaqaman, K. et al. Robust single-particle tracking in live-cell time-lapse sequences. Nat Meth 5, 695–702 (2008).

67. Sintorn, I.-M., Bischof, L., Jackway, P., Haggarty, S. & Buckley, M. Gradient based intensity normalization. J Microsc 240, 249–258 (2010).

