## Supplemental Materials for "Single molecule tracking reveals the role of transitory dynamics of nucleoid-associated protein HU in organizing the bacterial chromosome"

­Supplementary Figures

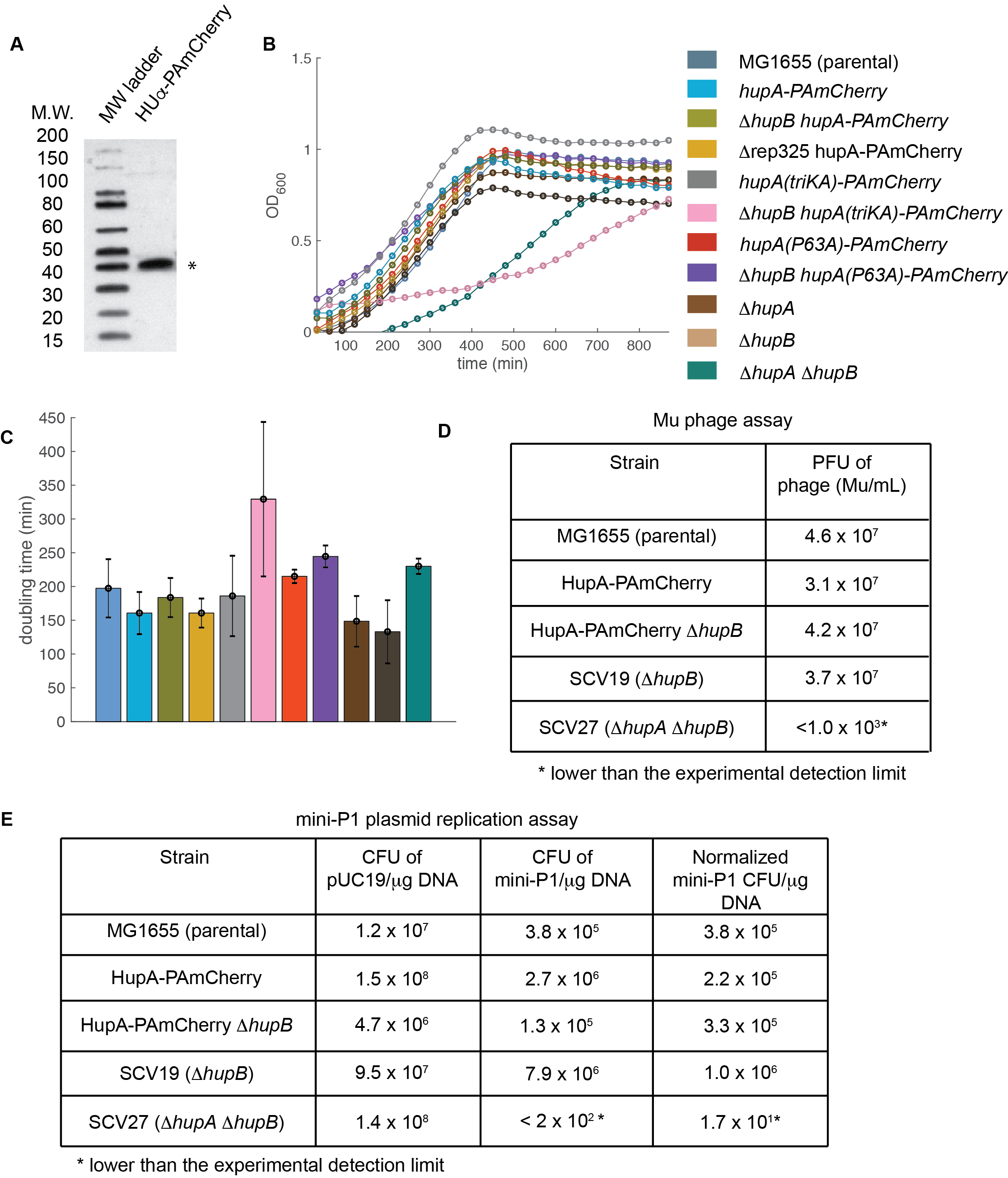

Supplementary Figure S1: HUα-PAmCherry fusion can replace the endogenous HUα for its function. (A) Western blot of Huα-PAmCherry in cells grown in rich growth media. Lane 1: protein ladder. Lane 2: cell lysate blotted against PAmCherry with * marking the correct molecular size of ~40 kDa for the fusion protein. (B) Representative growth curves of MG1655 (parental) and strains of different fusion and mutation backgrounds used in this study (Supplemental Table 1). (C) Averaged doubling times of all strains used in this study. Error bars represented standard deviation between at least three biological replicates. (D) Function of HUα-PAmCherry fusion measured by the ability of each strain to support the infection and plague formation of Mu phage. PFU indicates the phage plaque formation unit. (E) Function of HUα-PAmCherry fusion measured by its ability to support mini-P1 plasmid replication. The colony formation unit (CFU) after transformation of the mini-P1 plasmid into each strain was normalized against each strain's relative competency (CFU of a PUC19 plasmid transformation) compared to that of the wild type MG1655 parental strain.

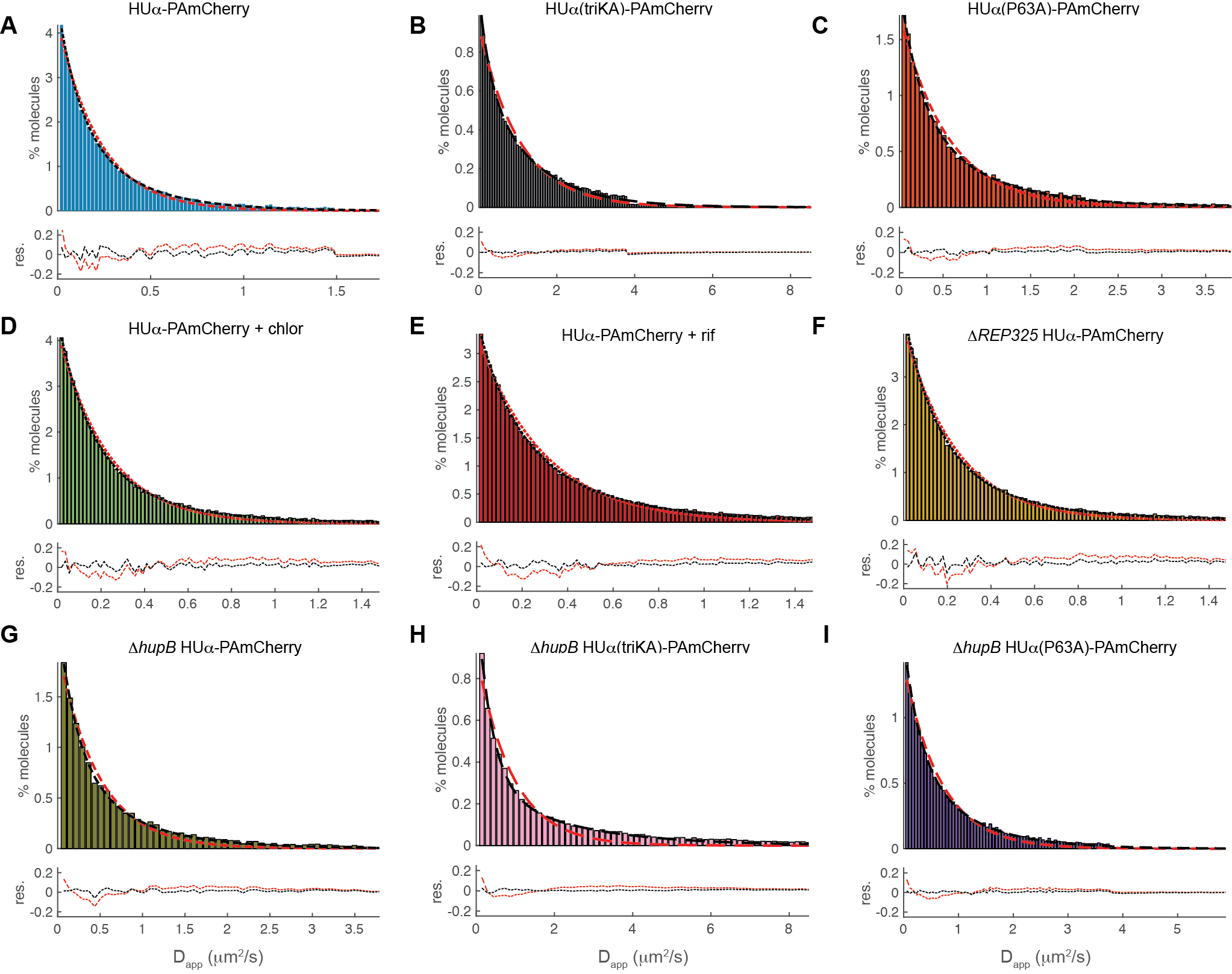

Supplemental Figure S2: Single-step diffusion coefficient distribution and the corresponding one-state (red) and two-state (black) fits (see Supplemental Note 1 for fitting details, and Supplemental Table 3 for fitting values) for HU-PAmCherry molecules in all strains and conditions used in the study. Residuals for the two different fits were shown below each distribution.

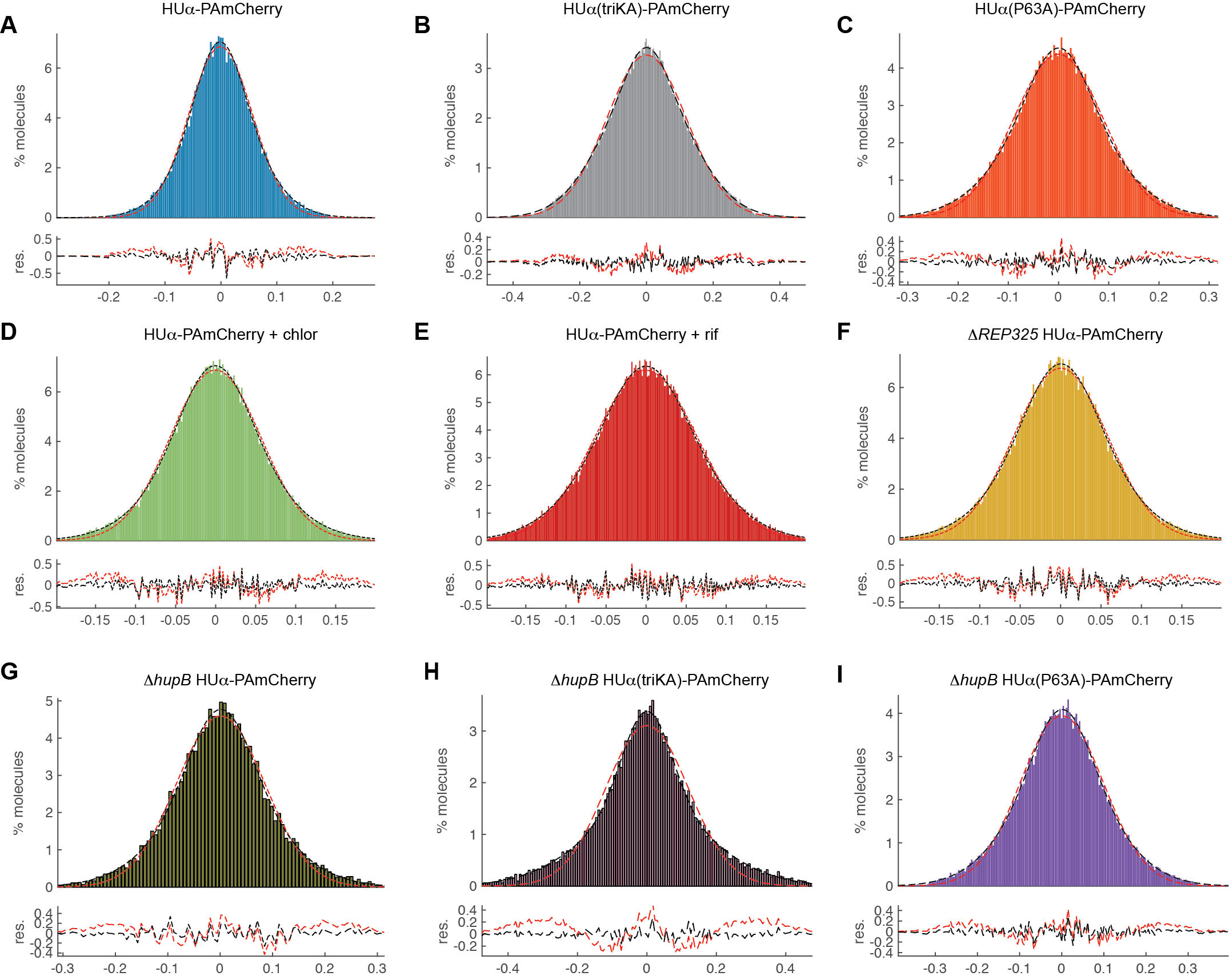

Supplemental Figure S3: Single-step displacement distribution along cell’s long-axis, and the corresponding one- (red) or two- (black) population Gaussian fits (see Supplemental Note 1 for fitting details, and Supplemental Table 3 for fitting values) for HU-PAmCherry molecules in all strains and conditions used in the study. Residuals for the two different fits were shown below each distribution.

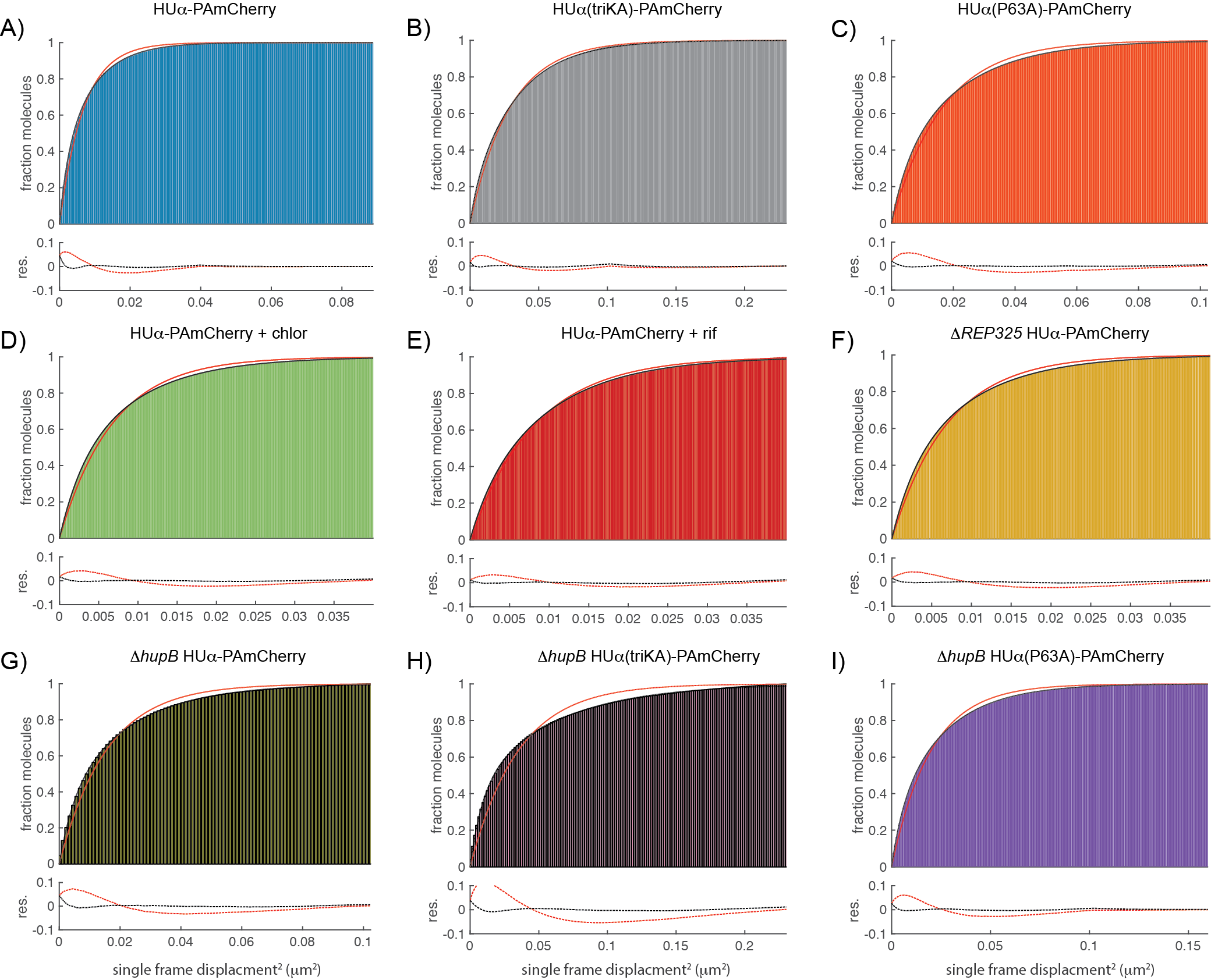

Supplementary Figure S4: Cumulative probability distribution calculated from single-step displacements and the corresponding one- (red) and two- (red) population fits (see Supplemental Note 1 for fitting details, and Supplemental Table 3 for fitting values) for HU-PAmCherry molecules in all strains and conditions used in the study. Residuals for the two different fits were shown below each distribution.

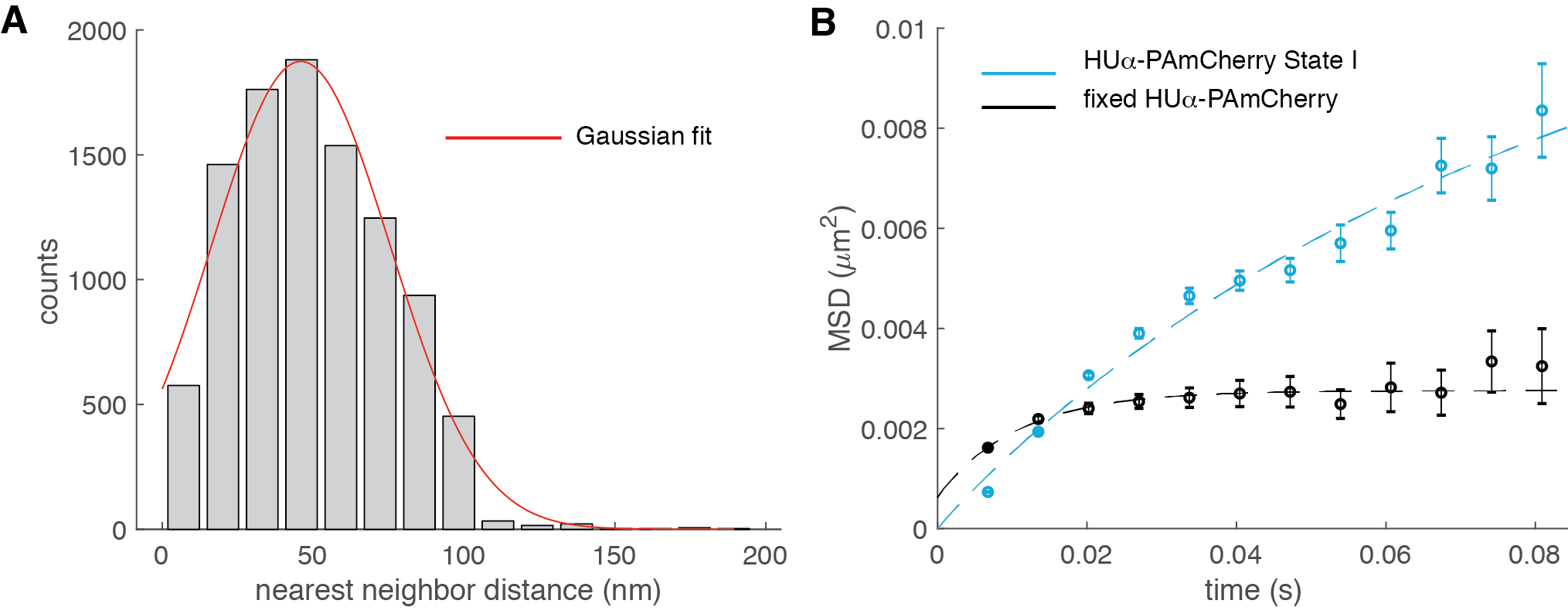

Supplementary Figure S5: Characterizations of the diffusion of HUα-PAmCherry molecules in fixed cells. (**A**) Nearest neighbor distance distribution of HUα-PAmCherry molecules was fit to a Gaussian function (red curve) to estimate the localization precision at ~ 45 nm (modified from^1^). (B) Apparent mean squared displacement plot of fixed HUα-PAmCherry molecules showed a plateau with an apparent diffusion domain of ~ 110 nm (dashed curve fit) using a confined diffusion model^2^, significantly smaller than that of State I HUα-PAmCherry molecules in live cells (blue curve).

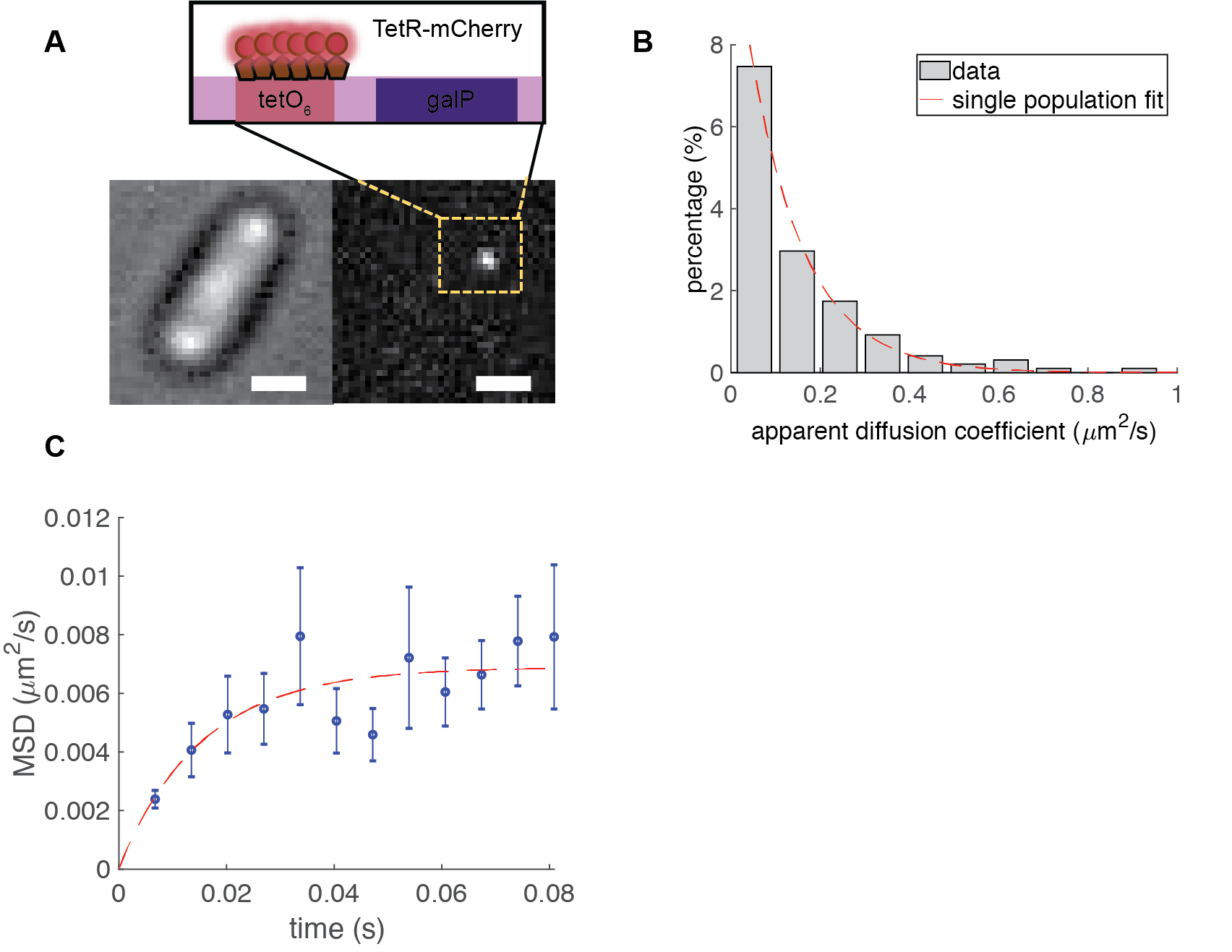

Supplemental Figure S6: Single molecule tracking of TetR-mCherry in a DNA fluorescent reporter operator (FROS) strain. (A) Schematic of FROS with representative fluorescent and brightfield images of a cell. Scale bar, 1 μm, (B) Distribution of apparent diffusion coefficients of individual TetR-mCherry molecules was best fit with a single-population (red dashed curve, *D*_app_ = 0.09 ± 0.04 μm^2^/s, μ ± s.e., n = 1808 displacements). (C) Mean squared displacement plot of TetR-mCherry trajectories with an apparent diffusive domain size of ~ 200 nm (dashed red curve). Error bars, standard error of the mean.

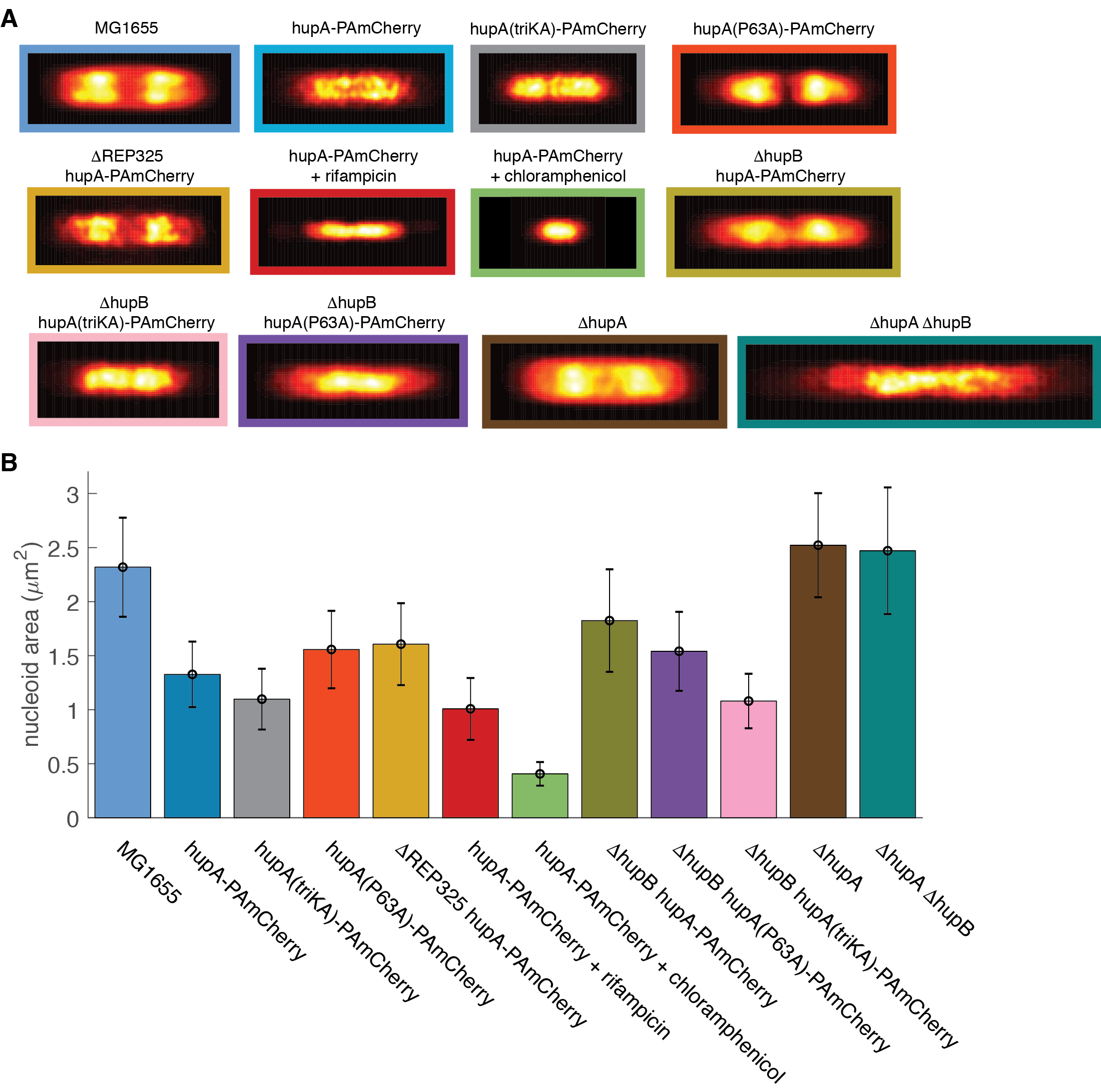

Supplementary Figure S7: SIM (structured illumination microscopy) images (**A**) and size quantifications (**B**) of nucleoids of all strains and conditions used in the study. (**A**) Normalized and averaged SIM images of nucleoids in all imaged conditions; Fluorescence intensity-normalized SIM images from at least twenty cells of each condition were rotated, aligned at cell centers and then superimposed with each other to generate the aggregated SIM image of that condition. (**B**) area of nucleoid at mid-plane for each strain and condition. Error bars, standard error of the mean calculated from bootstrapping.

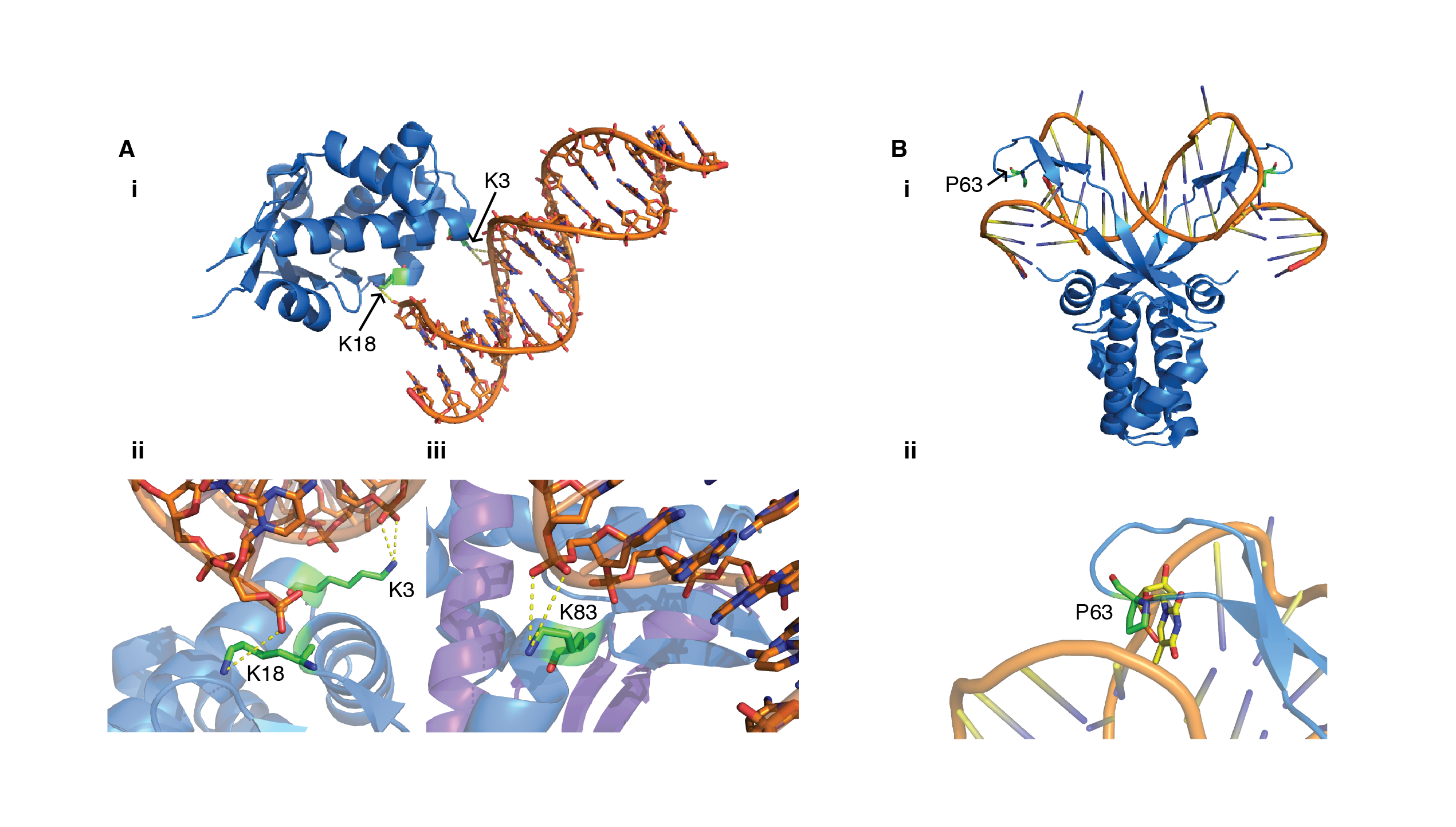

Supplementary Figure S8: Crystal structures of HU dimers in complex with nonspecific dsDNA (**A**) and pre-bent dsDNA (B). (Ai) Crystal structure of the *E. coli* HUα_2_ homodimer in complex with linear dsDNA (PDB: 4YEY) with a close-up view of lysine residues K3 and K18 in contact with the phosphate backbone of the DNA (Aii). (Aiii) Crystal structure of the *E. coli* HUαβ heterodimer in complex with linear dsDNA with the contact between DNA and lysine K83 residue highlighted (from PDB: 4FYT). (**Bi**) Crystal structure of Anabaena HU homodimer in complex with pre-bent DNA. The conserved proline residue intercalating into the DNA complex is highlighted and further magnified in (Bii).

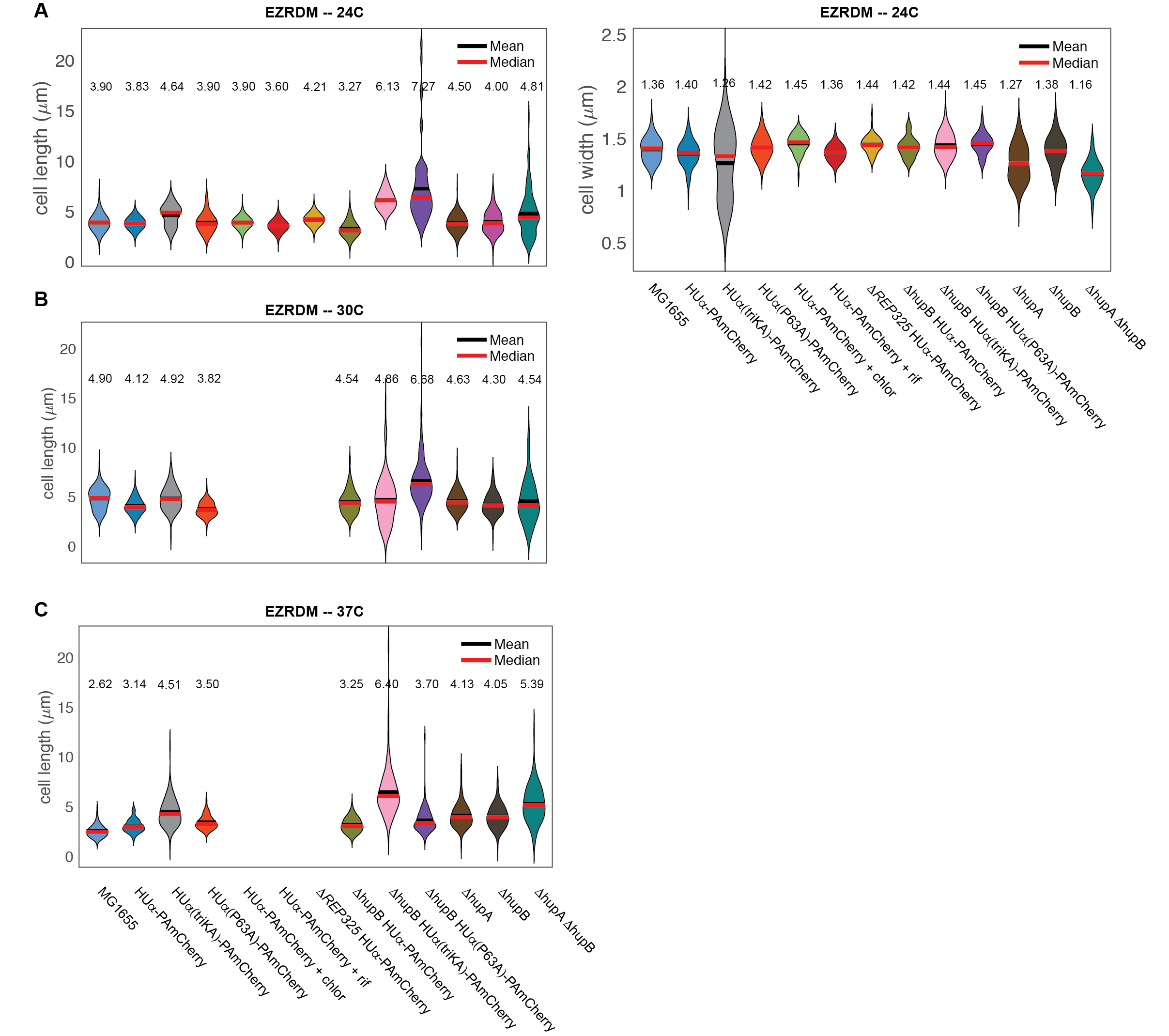

Supplemental Figure S9: Violin plots of cell sizes for all strains and controls grown in EZRDM at (**A**) 24°C, (**B**) 30°C, and (**C**) 37°C. Black bar, mean; red bars, median. All statistics (mean values, standard deviations, and cell numbers) for each strain were listed in Supplemental Table S4.

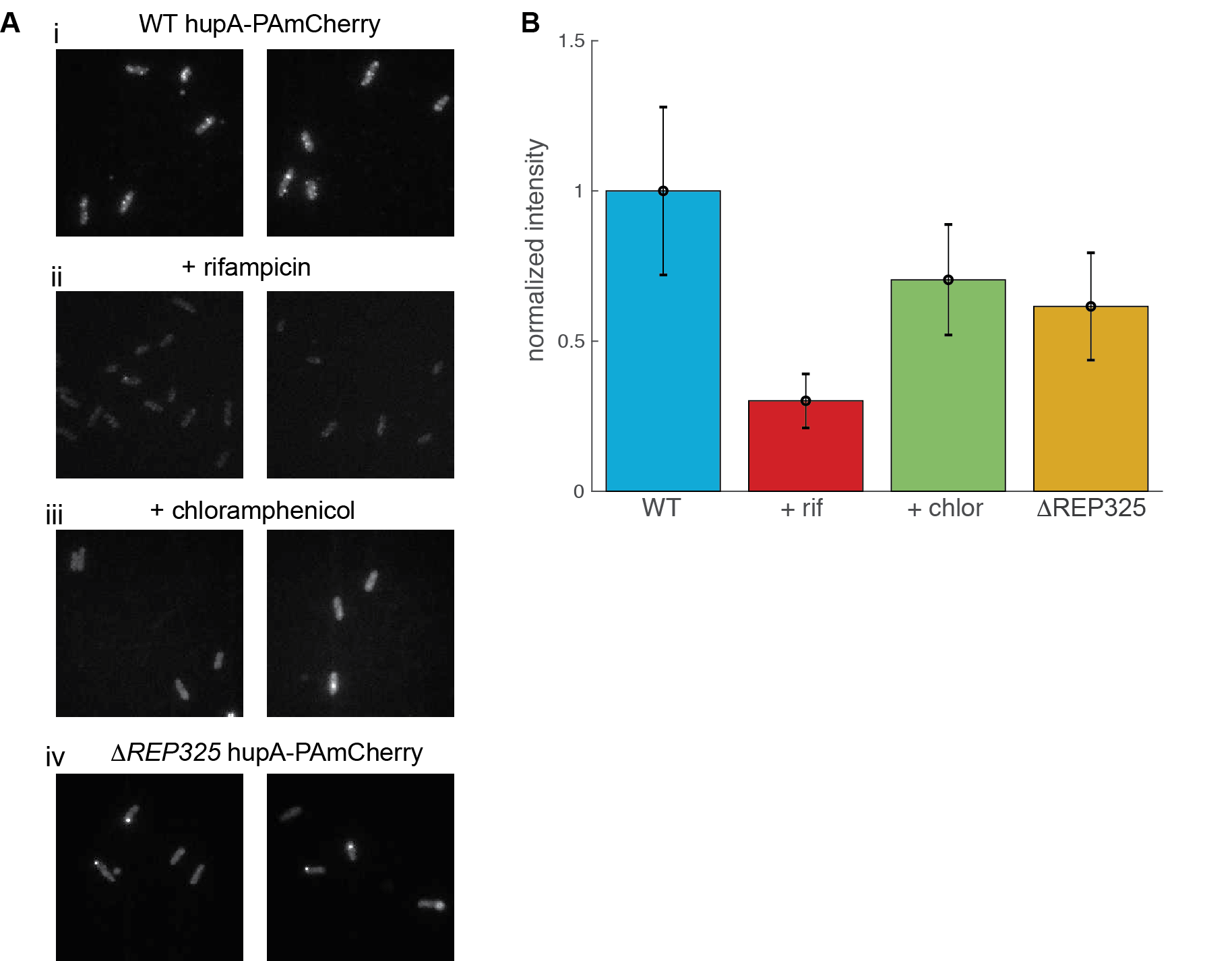

Supplementary figure S10: Single-molecule Fluorescence *in-situ* hybridization (smFISH) results of the ncRNA4 RNA under different conditions. (A) Example smFISH images of two biological replicate experiments for (i) WT *hupA-PAmCherry*, (ii) WT *hupA-PAmCherry* + rifampicin, (iii) WT *hupA-PAmCherry* + chloramphenicol, and (iv) Δ*REP325* *hupA-PAmCherry* cells. (B) Quantifications of smFISH signals by average intensity per cell; two replicates were pooled and normalized against the wild-type condition. Error bars represent combined standard deviation of individual cells and both biological replicates.

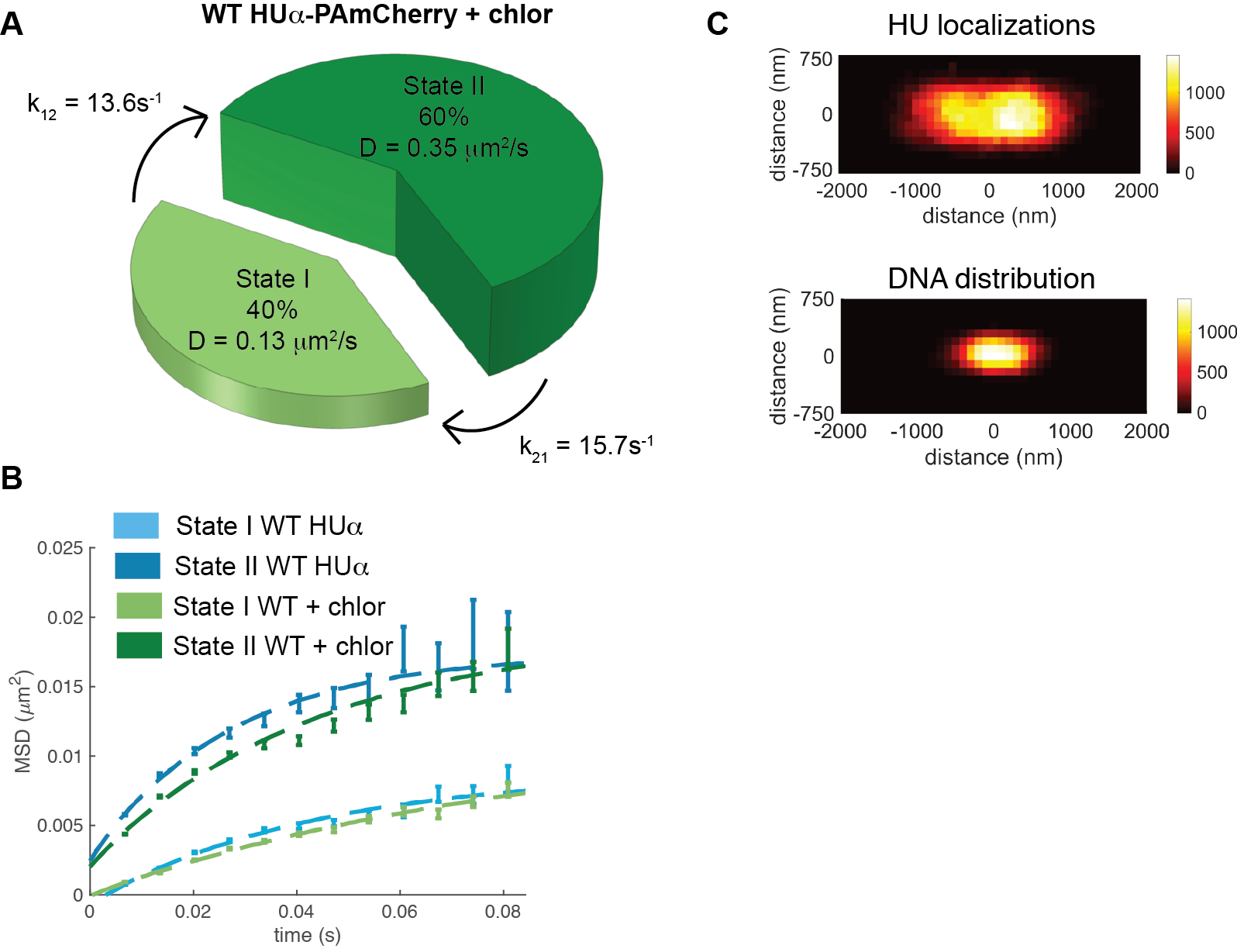

Supplementary Figure S11: HUα-PAmCherry diffusive dynamics and localizations in cells treated with chloramphenicol. (A) Two diffusive states of HUα-PAmCherry under chloramphenicol treatment with respective population percentages (size of pie piece), transition rates, and diffusion coefficients (height of pie piece) as identified by the HMM. (B) Mean squared displacement (MSD) plots of State I (light green) and State II (dark green) chlor-treated HUα-PAmCherry trajectories as a function of time compared to State I and II molecules in the WT condition (light and dark blue). (C) Two-dimensional (2D) histogram of all cellular chlor-treated HUα-PAmCherry localizations from SMT (top) and aggregated nucleoid morphology from SIM (structured illumination microscopy) imaging (bottom). The pixel size of both was 100 x 100 nm. The top color bar indicated the number of localizations used in each bin for HUα-PAmCherry (total 108,846 localizations). The bottom color bar indicated the normalized fluorescence level (in arbitrary units) of the nucleoid-intercalating dye Hoechst 33342 (total 20 fluorescence images).

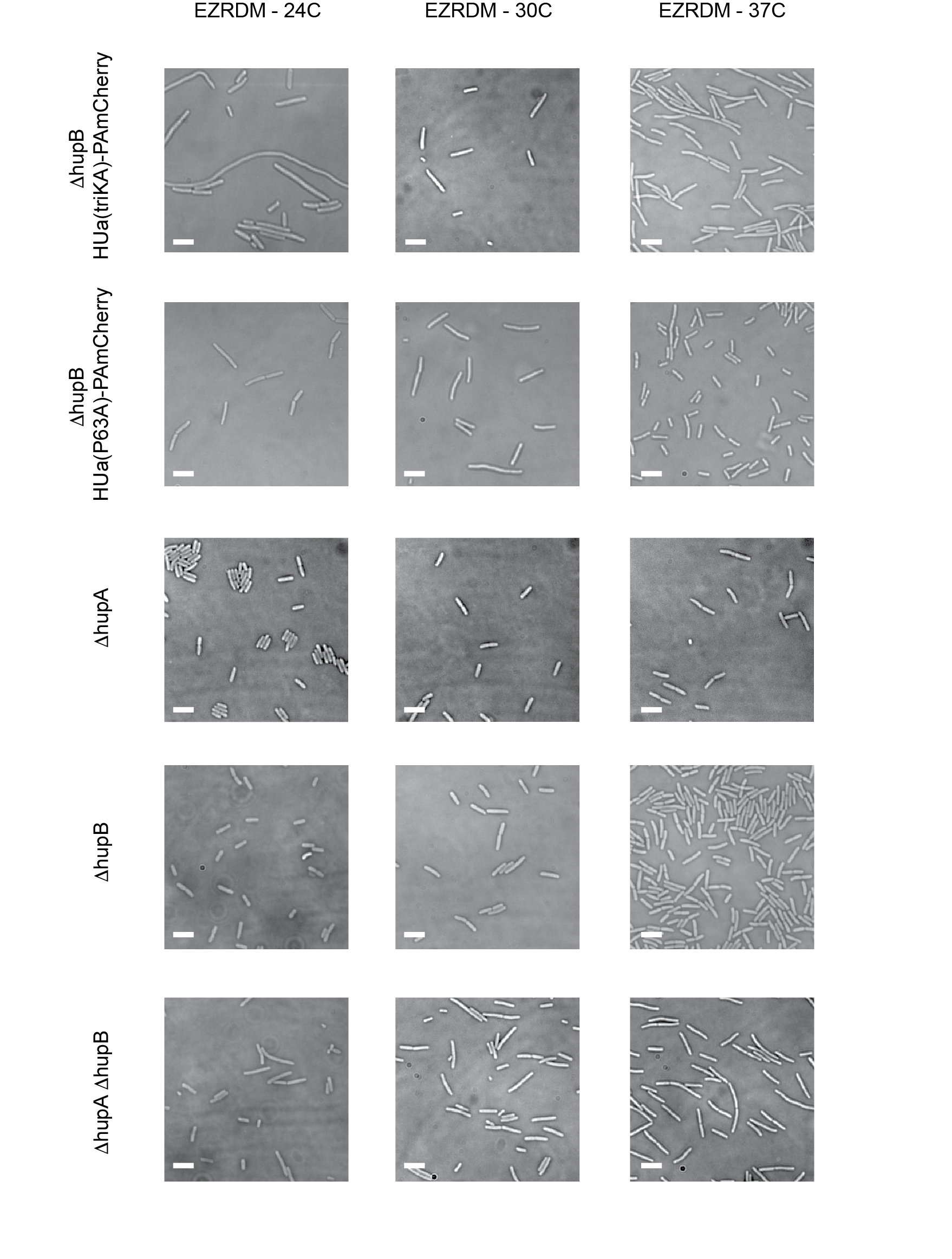

Supplementary Figure S12: bright-field imaging examples of deletion strains of HU. Each row represents a different deletion strain, each row represents the various temperatures cells were imaged: 24^o^C, 30^o^C, and 37^o^C. Scale bar in each image represents 5 microns.

Supplementary Table S1: Strains used in this study

| Strain | Genotype | Reference | Usage: |
| --- | --- | --- | --- |
| KB0026 | BW25113::hupA-PAmCherry-cmR | ***^3^*** | The “WT” hupA-PAmCherry strain used in Figure 1 |
| ACL035 | MG1655 galP::tetO_6_ pDR-tetR- mCherry | This study | For comparison to specific DNA-binding protein TetR |
| SCV148 | MG1655::hupA-PAmCherry-cmR | This study | For construction of Δ*hupB* *hupA-PAmCherry* (SCV153) |
| KB064 | KB0026 ΔREP325 | This study | Used in Figure 3ABC |
| SCV152 | SCV148 ΔhupB | This study | Δ*hupB hupA-PAmCherry* strain used in Figure 4ABC |
| SCV149 | MG1655::hupA(K3A K18A K83A)-PAmCherry | This study | Triple lysine mutant strain used in Figure 2ABC |
| SCV153 | SCV148 Δ*hupB* | This study | Triple lysine mutant strain in Δ*hupB* background used in Figure 4GHI |
| SCV146 | MG1655::hupA(P63A)-PAmCherry | This study | Proline mutant strain used in Figure 2DEF |
| SCV150 | SCV146 ΔhupB | This study | Proline mutant strain in Δ*hupB* background used in Figure 4DEF |
| SCV019 | MG1655 ΔhupA | This study | Δ*hupA* used for nucleoid volume comparisons |
| KB061 | MG1655 ΔhupB | Yale Genetic Stock | Δ*hupB* used for nucleoid volume comparisons and for construction of SCV 150, SCV152, and SCV153 |
| SCV020 | MG1655 ΔhupA ΔhupB | This study | Δ*hupA* Δ*hupB* used for nucleoid volume comparisons |

Supplementary Table S2: doubling times of all imaging and control strains.

| Strain | Replicate 1 | Replicate 2 | Replicate 3 |
| --- | --- | --- | --- |
| MG1655 (parental) | 159 min | 189 min | 244 min |
| HUα-PAmCherry | 149 min | 137 min | 196 min |
| ΔhupB HUα-PAmCherry | 170 min | 164 min | 217 min |
| ΔREP325 hupA-PAmCherry | 182 min | 139 min | 161 min |
| HUα(triKA)-PAmCherry | 189 min | 244 min | 125 min |
| ΔhupB HUα(triKA)-PAmCherry | 435 min | 345 min | 208 min |
| HUα(P63A)-PAmCherry | 208 min | 222 min | --- |
| ΔhupB HUα(P63A)-PAmCherry | 256 min | 233 min | --- |
| ΔhupA | 122 min | 175 min | --- |
| ΔhupB | 100 min | 166 min | --- |
| ΔhupA ΔhupB | 238 min | 222 min | --- |

Supplementary Table S3: diffusion coefficient values and occupation percentages for each diffusive states of HU-PAmCherry using various methodologies; 1D SFD is the single frame displacement distribution along the long axis of the cell, CDF is the cumulative probability distribution of all displacements, and 2D ADD is the apparent diffusion coefficient distribution in two dimensions; values reported are the mean values plus or minus the 95% confidence interval.

| *HUα-PAmCherry cells under rich growth media conditions* | | | | |
| --- | --- | --- | --- | --- |
| Methodology | D_1_ (μm^2^/s) | p_1_ (%) | D_2_ (μm^2^/s) | p_2_ (%) |
| 1D SFD | 0.14 ± 0.005 | 43 ± 5 | 0.38 ± 0.01 | 57 ± 6 |
| CDF | 0.12 ± 0.006 | 36 ± 3 | 0.35 ± 0.007 | 64 ± 3 |
| 2D ADD | 0.13 ± 0.07 | 26 ± 4 | 0.29 ± 0.08 | 74 ± 4 |
| HUα(triKA)-PAmCherry | | | | |
| Methodology | D_1_ (μm^2^/s) | p_1_ (%) | D_2_ (μm^2^/s) | p_2_ (%) |
| 1D SFD | 0.25 ± 0.08 | 11 ± 2 | 1.32 ± 0.08 | 89 ± 17 |
| CDF | 0.18 ± 0.03 | 12 ± 1 | 1.21 ± 0.03 | 88 ± 1 |
| 2D ADD | 0.28 ÷ 0.08 | 12 ± 0.1 | 1.30 ± 0.08 | 88 ± 0.1 |
| HUα(P63A)-PAmCherry | | | | |
| Methodology | D_1_ (μm^2^/s) | p_1_ (%) | D_2_ (μm^2^/s) | p_2_ (%) |
| 1D SFD | 0.26 ± 0.09 | 28 ± 3 | 0.89 ± 0.17 | 72 ± 10 |
| CDF | 0.19 ± 0.02 | 26 ± 2 | 0.79 ± 0.02 | 74 ± 4 |
| 2D ADD | 0.24 ± 0.06 | 26 ± 1 | 0.83 ± 0.1 | 74 ± 1 |
| *HUα-PAmCherry cells treated with chloramphenicol* | | | | |
| Methodology | D_1_ (μm^2^/s) | p_1_ (%) | D_2_ (μm^2^/s) | p_2_ (%) |
| 1D SFD | 0.14 ± 0.002 | 43 ± 2 | 0.4 ± 0.006 | 57 ± 3 |
| CDF | 0.11 ± 0.007 | 30 ± 3 | 0.32 ± 0.006 | 70 ± 3 |
| 2D ADD | 0.14 ± 0.04 | 37 ± 3 | 0.34 ± 0.08 | 63 ± 3 |
| *HUα-PAmCherry cells treated with rifampicin* | | | | |
| Methodology | D_1_ (μm^2^/s) | p_1_ (%) | D_2_ (μm^2^/s) | p_2_ (%) |
| 1D SFD | 0.15 ± 0.01 | 28 ± 5 | 0.42 ± 0.007 | 72 ± 5 |
| CDF | 0.12 ± 0.016 | 14 ± 2 | 0.35 ± 0.006 | 86 ± 2 |
| 2D ADD | 0.14 ± 0.06 | 22 ± 2 | 0.38 ± 0.07 | 78 ± 2 |
| *ΔREP325 HUα-PAmCherry cells* | | | | |
| Methodology | D_1_ (μm^2^/s) | p_1_ (%) | D_2_ (μm^2^/s) | p_2_ (%) |
| 1D SFD | 0.14 ± 0.05 | 38 ± 7 | 0.39 ± 0.12 | 62 ± 13 |
| CDF | 0.10 ±0.01 | 27 ± 3 | 0.33 ± 0.01 | 73 ± 3 |
| 2D ADD | 0.13 ± 0.05 | 29 ± 0.2 | 0.33 ± 0.07 | 71 ± 0.2 |
| *ΔhupB HUα-PAmCherry* | | | | |
| Methodology | D_1_ (μm^2^/s) | p_1_ (%) | D_2_ (μm^2^/s) | p_2_ (%) |
| 1D SFD | 0.26 ± 0.13 | 38 ± 6 | 0.92 ± 0.37 | 63 ± 9 |
| CDF | 0.17 ± 0.04 | 32 ± 5 | 0.81 ± 0.05 | 68 ± 5 |
| 2D ADD | 0.23 ± 0.09 | 32 ± 1 | 0.77 ± 0.17 | 68 ± 1 |
| *ΔhupB HUα(triKA)-PAmCherry cells* | | | | |
| Methodology | D_1_ (μm^2^/s) | p_1_ (%) | D_2_ (μm^2^/s) | p_2_ (%) |
| 1D SFD | 0.33 ± 0.03 | 40 ± 3 | 2.2 ± 0.08 | 59 ± 3 |
| CDF | 0.38 ± 0.07 | 35 ± 8 | 2.5 ± 0.04 | 65 ± 16 |
| 2D ADD | 0.38 ± 0.8 | 35 ± 0.01 | 2.23 ± 0.4 | 65 ± 0.01 |
| *ΔhupB HUα(P63A)-PAmCherry cells* | | | | |
| Methodology | D_1_ (μm^2^/s) | p_1_ (%) | D_2_ (μm^2^/s) | p_2_ (%) |
| 1D SFD | 0.2 ± 0.02 | 25 ± 2 | 0.94 ± 0.002 | 75 ± 2 |
| CDF | 0.3 ± 0.1 | 29 ± 2 | 1.1 ± 0.2 | 71 ± 5 |
| 2D ADD | 0.29 ± 0.06 | 25 ± 0.01 | 1.02 ± 0.1 | 75 ± 0.01 |

Supplementary Table S4: cell lengths and width for imaging and control strains in EZRDM and LB medias under increasing temperatures (24^o^C, 30^o^C, and 37^o^C). Number of cells (N) for each condition are listed after the mean value in parentheses. Error listed is the standard deviation of the sample.

| Strain | Cell length – EZRDM 24^o^C (μm) | Cell width – EZRDM 24^o^C (μm) | Cell length – EZRDM 30^o^C (μm) | Cell length – EZRDM 37^o^C (μm) |
| --- | --- | --- | --- | --- |
| MG1655 (parental) | 3.9 ± 0.8 (N=109) | 1.4 ± 0.1 (N=100) | 5.0 ± 1.1 (N=131) | 2.6 ± 0.6 (N=128) |
| HUα-PAmCherry | 3.8 ± 0.4 (N=75) | 1.4 ± 0.1 (N=75) | 4.1 ± 0.8 (N=95) | 3.1 ± 0.6 (N=157) |
| HUα(triKA)-PAmCherry | 4.6 ± 1.0 (N=61) | 1.3 ± 0.3 (N=61) | 4.9 ± 1.2 (N=102) | 4.5 ± 1.2 (N=102) |
| HUα(P63A)-PAmCherry | 3.9 ± 0.9 (N=140) | 1.4 ± 0.1 (N=60) | 3.8 ± 0.7 (N=102) | 3.5 ± 0.7 (N=152) |
| HUα-PAmCherry + chlor. | 3.9 ± 0.4 (N=61) | 1.5 ± 0.1 (N=61) |  |  |
| HUα-PAmCherry + rif. | 3.6 ± 0.4 (N=87) | 1.4 ± 0.1 (N=87) |  |  |
| Δ*REP325* HUα-PAmCherry | 4.2 ± 0.4 (N=78) | 1.4 ± 0.1 (N=78) |  |  |
| Δ*hupB* HUα-PAmCherry | 3.3 ± 0.7 (N=150) | 1.4 ± 0.1 (N=48) | 4.5±1.2 (N=102) | 3.3 ± 0.7 (N=121) |
| Δ*hupB* HUα(triKA)-PAMCherry | 6.1 ± 0.9 (N=67) | 1.4 ± 0.1 (N=68) | 4.8 ± 2.4 (N=109) | 6.5 ± 2.4 (N=108) |
| Δ*hupB* HUα(P63A)-PAmCherry | 7.3 ± 3.6 (N=71) | 1.5 ± 0.1 (N=67) | 6.7 ± 2.4 (N=133) | 3.7 ± 1.1 (N=159) |
| Δ*hupA* | 3.8 ± 0.8 (N=121) | 1.3 ± 0.2 (N=100) | 4.6 ± 1.0 (N=149) | 4.2 ± 1.2 (N=128) |
| Δ*hupB* | 4.0 ± 1.0 (N=109) | 1.4 ± 0.1 (N=95) | 4.3 ± 1.1 (N=104) | 4.0 ± 1.1 (N=104) |
| Δ*hupA* Δ*hupB* | 4.8 ± 2.3 (N=94) | 1.2 ± 0.1 (N=101) | 4.7 ± 2.2 (N=110) | 5.4 ± 1.7 (N=103) |

| Supplementary Table S5: confinement values (as calculated using the equation in Supplemental Note 2) for each state of HUα-PAmCherry in drug treatments and mutant strains. Values listed are μ ± std from bootstrapping. | | | | |
| --- | --- | --- | --- | --- |
| Strain |  | State 1 (nm) |  | State 2 (nm) |
| HUα-PAmCherry |  | 239 ± 1 |  | 299 ± 1 |
| HUα(triKA)-PAmCherry |  | 345 ± 1 |  | n/a |
| HUα(P63A)-PAmCherry |  | 942 ± 7 |  | 796 ± 1 |
| HUα-PAmCherry + chlor |  | 272 ± 1 |  | 320 ± 1 |
| HUα-PAmCherry + rif |  | n/a |  | 366 ± 1 |
| Δ*REP325* HUα-PAmCherry |  | 392 ± 2 |  | 452 ± 1 |
| HUα-PAmCherry + Δ*hupB* |  | 428 ± 2 |  | n/a |
| HUα(triKA)-PAmCherry + Δ*hupB* |  | 632 ± 1 |  | 1344 ± 1 |
| HUα(P63A)-PAmCherry + Δ*hupB* |  | 849 ± 4 |  | 1099 ± 1 |

Supplementary Table S6: estimated K_d_ of each HU mutant and growth condition to dsDNAs using the transition probabilities from HMM (tabulated in Table 1). K_d_ measurements are calculated as described in Supplementary Note 3.

|  | k_21_ (s^-1^) | k_12_ (s^-1^): off rate | on time (ms, State I) | k_on_ (M^-1^s^-1^) on rate | off time (ms, State II) | K_d_ (M) |
| --- | --- | --- | --- | --- | --- | --- |
| hupA-PAmCherry | 14.7 | 10.9 | 100 | 2.3E+03 | 74 | 4.8E-03 |
| hupA(triKA)-PAmCherry | 1.2 | 9.7 | 107 | 1.8E+02 | 852 | 5.2E-02 |
| hupA(P63A)-PAmCherry | 9.3 | 8.7 | 129 | 1.4E+03 | 114 | 6.1E-03 |
| hupA-PAmCherry + chlor | 15.7 | 13.6 | 80 | 2.4E+03 | 70 | 5.6E-03 |
| ΔREP325 hupA-PAmCherry | 12.9 | 13.8 | 72 | 2.0E+03 | 96 | 6.9E-03 |
| hupA-PAmCherry + rif | 10.3 | 12.4 | 80 | 1.6E+03 | 100 | 7.8E-03 |
| ΔHUβ hupA-PAmcherry | 6.8 | 6.2 | 168 | 1.0E+03 | 155 | 5.9E-03 |
| ΔHUβ hupA(P63A)-PAmCherry | 9.7 | 10.7 | 100 | 1.5E+03 | 110 | 7.2E-03 |
| ΔHUβ hupA(triKA)-PAmCherry | 12.6 | 5.4 | 196 | 1.9E+03 | 84 | 2.8E-03 |
| DNA bp | volume (L) | DNA concentration | |  |  |  |
| 9.20E+06 | 2.35E-15 | 6.49E-03 | |  |  |  |

Supplementary Table S7: top table is the cross-comparison of nucleoid areas measured from the SIM images as demonstrated in Figure S7; each number represents the p-value from comparing the row to the column calculated from performing a one-way ANOVA test. Bottom table is the statistics table from the one-way ANOVA test.

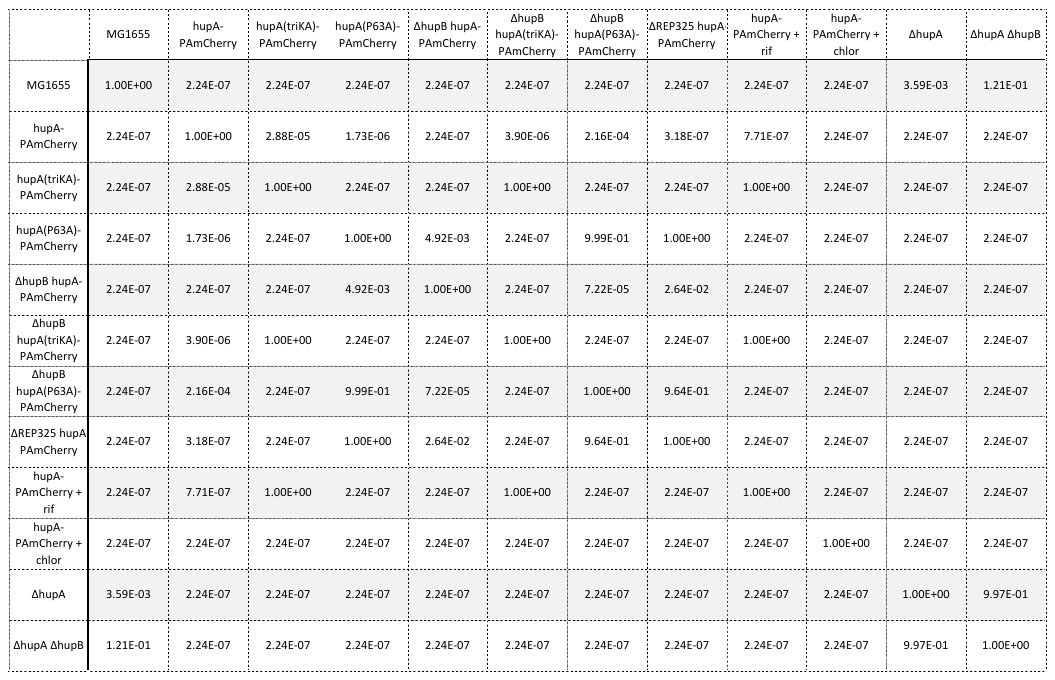

|  | SS | df | MS | F | Prob > F |
| --- | --- | --- | --- | --- | --- |
| Columns | 450.14 | 11.00 | 40.92 | 330.33 | 0.00 |
| Error | 147.17 | 1188.00 | 0.12 |  |  |
| Total | 597.31 | 1199.00 |  |  |  |

Supplementary Table S8: effects of various perturbations of HU on various cell phenotypes, including cell growth, cell length, condensation, and segregation. A single (+) indicates a normal phenotype; (++) indicates increased growth/length/nucleoid size while (-) or (--) indicates decreased growth/length/nucleoid size. For segregation, (+) indicates a visible while (-) indicates the lack of discernable central cleft between the two nucleoids normally present in fast-growing cells.

| Strain | Cell growth | Cell Length | Conden-sation | Segre-gation |
| --- | --- | --- | --- | --- |
| MG1655 (parental) | **+** | **+** | **+** | **+** |
| HUα-PAmCherry | **+** | **+** | **+** | **+** |
| ΔhupB HUα-PAmCherry | **+** | **+** | **-** | **+** |
| ΔREP325 hupA-PAmCherry | **+** | **+** | **+** | **+** |
| HUα(triKA)-PAmCherry | **+** | **-** | **-** | **-** |
| ΔhupB HUα(triKA)-PAmCherry | **--** | **++** | **-** | **-** |
| HUα(P63A)-PAmCherry | **+** | **+** | **++** | **+** |
| ΔhupB HUα(P63A)-PAmCherry | **--** | **++** | **++** | **+** |
| ΔhupA | **+** | **+** | **-** | **+** |
| ΔhupB | **+** | **+** | **-** | **+** |
| ΔhupA ΔhupB | **--** | **++** | **--** | **-** |

Supplementary Table S9: Total cellular RNA content from equivalent O.D. of cells under different conditions.

| Condition | Total RNA | Quality *A*_260_/*A*_280_ | Normalized to WT |
| --- | --- | --- | --- |
| WT | 35065 ng | 1.94 | 1 |
| Rifampicin (60 min) | 15655 ng | 1.91 | 0.45 |
| Chloramphenicol (60 min) | 35755 ng | 1.97 | 1.02 |

**Supplemental Information**

**Supplemental Note 1: Calculation of diffusion coefficients**

Several methods were employed to calculate the best number of diffusion coefficients represented in the data and their diffusion coefficients. First, we calculated all single frame displacements along the x-axis. To achieve this, we manually rotated all cells such that the long axis of the cell was defined as the x-axis and rotated all the trajectories by the same angle. We chose to take displacements along the x-axis as there is less likelihood of these particles encountering the barrier of the cell wall^4^. We found the single-frame displacement along the x-axis and plotted the histogram of their distribution binned into a number of bins equal to the square root of total number of displacements. This distribution resembles a Gaussian centered around zero; due to the nature of Brownian motion, displacements on average will be zero and the width of the distribution is related to the number of diffusion coefficients represented in the data and their values. We fit this distribution of single-frame displacements to the following equation:

$$p\left( r \right)=\sum_{i=1}^{n} \frac{p_{i}}{\sqrt{4\pi D_{i}t}}e^{\frac{-r^{2}}{4D_{i}t}}$$

Where *r* is the single frame displacement, *n* is the total number of diffusive states represented in the data, *D_i_* is the diffusion coefficient of the *i*th diffusive state, and *p_i_* is the fraction of molecules in the *i*th state. To determine the best number of states, we fit our data to models for one, two, and three diffusive states. We saw no improvement of the fit and dispersion of residuals when we moved from a two-state to a three-state model, so we deemed the two-state model to be the best fit.

Secondly, we calculated the cumulative displacement probability distribution. Similar to above, we calculated the displacement between consecutive frames for each trajectory. To estimate the cumulative displacement probability distribution, we divided the displacements into a number of bins equal to the squre root of the total number of displacements. For each displacement bin, we calculated the fraction of displacements less than or equal to the bin value. This cumulative displacement probability distribution can be fit to the following equation:

$$cdf\left( r \right)=1- \sum_{i=1}^{n} p_{i}e^{\frac{{-x}^{2}}{4D_{i}t}}$$

Where *r* is the single frame displacement, *n* is the total number of diffusive states represented in the data, D*_i_* is the diffusion coefficient of the *i*th diffusive state, and p*_i_* is the fraction of molecules in the *i*th state such that the sum of all pi values equals one. To determine the best number of states, we fit our data to one, two, and three-state models. We saw no improvement of the fit and dispersion of residuals when we moved from a two-state to a three-state model, so we deemed the two-state model to be the best fit.

Lastly, we calculated the apparent diffusion coefficient from each individual displacement. We binned the apparent diffusion coefficients into twelve bins and plotted their distribution. This distribution of apparent diffusion coefficients can be fit to the equation:

$$p\left( D \right)= \sum_{i=1}^{n} \frac{a_{i}}{D_{i}}e^{-\frac{1}{D_{i}}}$$

where *n* is the number of diffusive states represented in the data, *a_i_* is the fraction of molecules in the *i*th diffusive state, and *D_i_* is the diffusion coefficient of the ith diffusive state. Again, we determined the best number of state by fitting our data to one, two, and three-state models. We saw no improvement of the fit when we moved from a two-state to a three-state model, so again we deemed the two-state model to be the best fit.

**Supplemental Note 2: Calculation of confinement zone from MSD data**

First, we calculated the mean-squared displacement (MSD) for 15 timelags by using the following equation:

$$MSD\left( t \right)=\frac{1}{N}\sum_{n=1}^{N} {(x_{n}\left( t \right)-x_{n}\left( 0 \right))}^{2}$$

Where N is the total number of displacements to be averaged for that timelag, *x_n_*(0) is the initial coordinates of each trajectory, and *x_n_*(t) is the coordinates at time t. For pure Brownian motion, this normally is linear such that MSD(t) = 4Dt; i.e. the slope of the line is proportional to the diffusion coefficient of the molecule. MSD can be thought of as the area that a collection of particles covers in time. Thus, if particles encounter a barrier, then as time approaches infinity, the MSD remains unchanged. Kusumi, et. al. derived the equation for diffusion within a boundary of length *L* ^2^, which is:

$$MSD(t)= L^{6}+\frac{{16 L}^{2}}{\pi^{4}}\sum_{i=1}^{\infty} e^{\frac{-\left( i\pi\sigma\right)^{2}}{L^{2}t}}$$

where *L* is the total width of the boundary, and σ is the average displacement such that D = σ^2^/2. We approximated the sum up to one hundred steps, ensuring we reached convergence, and used non-linear least squares fitting in MATLAB to find the parameters that gave the least squared residuals. To estimate the error in the fitting parameters, we performed a pseudo-bootstrapping method in which we removed a random trajectory from our data set and estimated the parameters again. We repeated this pseudo-bootstrapping one hundred times for each curve; the error reported was the standard deviation of the values from these bootstrapping calculations.

**Supplemental Note 3: Calculation of the rate constants and dissociation constant for HU**

Our HMM of HU gives transition probabilities that can be converted into kinetic rate constants. Given a transition matrix A, we can convert these transition probabilities into kinetic rate constants through this relation:

$$A=\left[ \begin{matrix} a_{ij} & \cdots& ain \\ \vdots& \ddots& \vdots\\ anj & \cdots& ann \end{matrix} \right]=exp\left( \left[ \begin{matrix} kij & \cdots& kin \\ \vdots& \ddots& \vdots\\ knj & \cdots& knn \end{matrix} \right]\cdot\Delta t \right)$$

where *a_ij_* is the transition probability of a molecule transitioning from state *i* to state *j* within the frame rate *Δt*, and *k_ij_* is the corresponding kinetic rate between the *i*th and *j*th diffusive state. We assigned state I HU-PAmCherry molecules to be bound to the DNA, and state II HU-PAmCherry molecules to be loosely associated, we can assume that HU binds to the DNA using this reaction:

$$HU+DNA \begin{matrix} \underset{\leftarrow}{k_{off}} \\ \overset{\to}{k_{on}} \end{matrix} HU\cdot DNA$$

Assuming that [DNA] >> [HU], we can use a first order approximation, where K_d_ = k_off_[DNA}/k_on_. We assume that each basepair of the DNA provides a unique binding site for the DNA and assume that on average, there are two full chromosomes per *E. coli* cell under our fast growth conditions. Converting the total number of basepairs into molar concentration gives a concentration of DNA around 4mM. Because we know the k_on_ and k_off_ values from our HMM, we can then estimate the dissociation rate for each condition.

Supplemental References:

1.	Endesfelder, U., Malkusch, S., Fricke, F. & Heilemann, M. A simple method to estimate the average localization precision of a single-molecule localization microscopy experiment. *Histochem. Cell Biol.* **141,** 629–638 (2014).

2.	Kusumi, A., Sako, Y. & Yamamoto, M. Confined lateral diffusion of membrane receptors as studied by single particle tracking (nanovid microscopy). Effects of calcium-induced differentiation in cultured epithelial cells. *Biophysical Journal* **65,** 2021–2040 (1993).

3.	Wang, S., Moffitt, J. R., Dempsey, G. T., Xie, X. S. & Zhuang, X. Characterization and development of photoactivatable fluorescent proteins for single-molecule-based superresolution imaging. *Proc. Natl. Acad. Sci. U.S.A.* **111,** 8452–8457 (2014).

4.	Bohrer, C. H., Bettridge, K. & Xiao, J. Reduction of Confinement Error in Single-Molecule Tracking in Live Bacterial Cells Using SPICER. *Biophysical Journal* **112,** 568–574 (2017).
